# Cilia Directionality Reveals a Slow Reverse Movement of Principal Neurons for Positioning and Lamina Refinement in the Cerebral Cortex

**DOI:** 10.1101/2021.12.21.473383

**Authors:** Juan Yang, Soheila Mirhosseiniardakani, Liyan Qiu, Kostandina Bicja, Abigail Del Greco, Kevin Jungkai Lin, Mark Lyon, Xuanmao Chen

## Abstract

Currently little is known about neuronal positioning and the roles of primary cilia in postnatal neurodevelopment. We show that primary cilia of principal neurons undergo marked changes in positioning and orientation, concurrent with postnatal neuron positioning in the mouse cerebral cortex. Primary cilia of early- and late-born principal neurons in compact layers display opposite orientations, while neuronal primary cilia in loose laminae are predominantly oriented toward the pia. In contrast, astrocytes and interneurons, and neurons in nucleated brain regions do not display specific cilia directionality. We further discovered that the cell bodies of principal neurons in inside-out laminated regions spanning from the hippocampal CA1 region to neocortex undergo a slow “reverse movement” for postnatal positioning and lamina refinement. Further, selective disruption of cilia function in the forebrain leads to altered lamination and gyrification in the retrosplenial cortex (RSC) that is formed by reverse movement. Collectively, this study identifies reverse movement as a fundamental process for principal cell positioning that refines lamination in the cerebral cortex and casts light on the evolutionary transition from 3-layered allocortices to 6-layered neocortices.

**Summary Statement:** Guided by cilia directionality, we discovered that the somata of principal neurons in inside-out laminated regions undergo a slow but substantial reverse movement for postnatal cell positioning, which refines cortical layering.

## Introduction

Primary cilia are centriole-derived organelles that exists in almost all mammalian cell types, including neurons (Gopalakrishnan et al., 2023; Jurisch-Yaksi et al., 2024; Mill et al., 2023; Park et al., 2019). Primary cilia mediate numerous function and their defects are associated with a variety of disorders including neurodevelopmental disorders (Anvarian et al., 2019; Hilgendorf et al., 2024; Kumar and Reiter, 2021; Lee and Gleeson, 2010; Park et al., 2019; Reiter and Leroux, 2017; Singla and Reiter, 2006). It is well-known that primary cilia modulate embryonic neurodevelopment by influencing cell fate specification in the ventral neural tube (Mukhopadhyay et al., 2013; Pal and Mukhopadhyay, 2015) or forebrain neuronal patterning (Dessaud et al., 2008; Gorivodsky et al., 2009). However, how cilia regulate postnatal neurodevelopment remains poorly understood.

The cerebral cortex is composed of the neocortex, the allocortex and neighboring structures (Aboitiz and Zamorano, 2013). The neocortex contains 6 sparse layers, with principal (projection or excitatory) neurons distributed across Layers II-VI (Klingler, 2017). The allocortex, such as the mouse hippocampus, consists of three laminae (Laurent et al., 2016; Luzzati, 2015), where most principal neurons are compacted into Layer II. Thus far, it is unclear how the 6-layered neocortex evolved from the 3-layered allocortex (Luzzati, 2015; Rakic, 2009). Cortical laminae of principal neurons are divided into the deep and superficial sublayers along the radial axis (Cembrowski and Spruston, 2019; Soltesz and Losonczy, 2018). These two sublayers harbor early- and late-born principal neurons, respectively, which migrate to destination in a birthtime-dependent inside-out pattern (Khalaf-Nazzal and Francis, 2013; Soltesz and Losonczy, 2018). Following the radial migration (Kriegstein and Noctor, 2004; Yoshinaga et al., 2021), principal neurons are subject to positioning and subsequently become matured (Liu et al., 2019; Moffat et al., 2015). The first two weeks after birth represent the critical period for neuronal positioning, when the hippocampal superficial and deep sublayers gradually condense into the compact stratum pyramidale (SP), whereas neocortical principal neurons expands to form five sparse laminae (Layers II-VI) (Geschwind and Rakic, 2013). It remains elusive how principal neurons in the cerebral cortex complete positioning during postnatal development.

A recent studies have shown that primary cilia regulate the migration of neuroblast through the rostral migratory stream (RMS) and the orientation of primary cilia is linked to microtubules dynamics when neurons stay at different migrating phases (Matsumoto et al., 2019). Because the centrosome modulates microtubule network dynamics that, in turn, influences the position of the nucleus in neurons during neurodevelopment (Higginbotham and Gleeson, 2007; Kuijpers and Hoogenraad, 2011; Lee and Gleeson, 2011; Tsai and Gleeson, 2005), cilia emanation out of centrioles and cilia orientation may provide useful clues to understand the process of neuron positioning. We first performed cilia morphology and directionality analyses in the mouse postnatal cerebral cortex. We observed that primary cilia of principal neurons display specific directionality. This clue gradually led us to discover that principal neurons in inside-out laminated regions undergo a slow reverse movement for positioning. Further, selective cilia ablation in the forebrain causes altered cortical laminae and gyrification in RSC that is formed by backward movement. Together, we have identified reverse movement as a fundamental process for principal neuron positioning, crucial not only for the postnatal development of the cerebral cortex, but also for elucidating the evolutionary transition from the 3-layered allocortices to the 6-layered neocortices.

## Results

### Primary Cilia of Early- and Late-born CA1 Neurons Display Opposite Orientations

To examine cilia morphology changes in CA1 SP during postnatal development, we performed immunofluorescence staining on WT brain sections using adenylyl cyclase 3 (AC3) (Bishop et al., 2007; Chen et al., 2016; Yang et al., 2020; Zhou et al., 2019) and ADP-ribosylation factor-like protein 13B (ARL13B) antibodies (Sterpka and Chen, 2018; Sterpka et al., 2020). We found that cilia morphology and orientation in SP underwent marked changes postnatally (Fig. 1A-D). At postnatal day 5 (P5) and P7, AC3- or ARL13B-marked cilia were rarely detected in CA1 SP (Fig. 1A). At P9, many neuronal cilia became visible but still very short, and at P10, cilia were markedly elongated, with many clustered at the edge of SP and stratum radiatum (SR) (Fig. 1D). Clearly, cilia exhibited opposite orientations: cilia axoneme in the superficial sublayer mostly were oriented to SR, whereas those in the deep sublayer oriented towards the stratum oriens (SO). Interestingly, from P10 to P14, cilia orientation had a drastic change (Fig. 1D). At P10, ∼ 40% of cilia oriented toward SO, while at P14 only ∼20% of cilia maintained that orientation (Fig. 1A-E). At P14, SR-oriented cilia occupied most of SP space. Afterwards at P40 and P75, cilia orientation did not change much, except for becoming less uniformly aligned (Fig. 1A-E). Simultaneously, the thickness of SP decreased significantly (Fig. 1F) and continued to decrease with age (Fig. S1A). These data suggest that pyramidal neurons continue to adjust their soma position postnatally, decreasing SP lamina thickness when some neurons die (Wong and Marin, 2019).

**Fig. 1.**
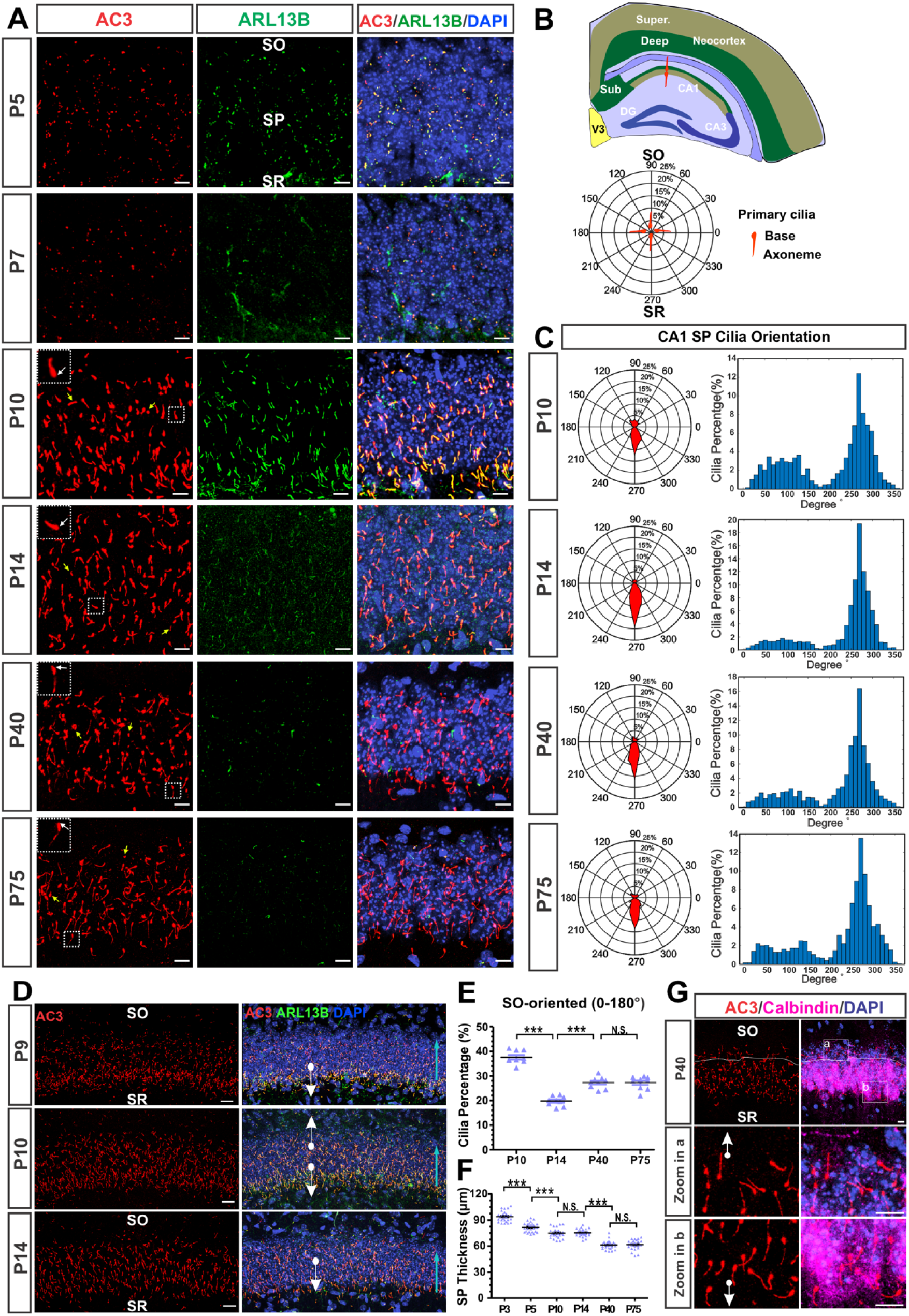
Opposite primary cilia orientation in CA1 SP of WT mice. **(A)** SP neuronal cilia images at different ages. White arrows indicate the base of the cilia. Yellow arrow points to cilia that were unquantifiable in terms of orientation. Scale bars: 10 µm. **(B)** A diagram showing cilia in the dorsal hippocampal CA1 SP. Cilia are posited in the polar plot. The numbers (0-360°) on the out circles denote cilia orientation. **(C)** The percentage of cilia axonemes oriented toward SO (0 -180°) and SR (180 - 360°) is shown in polar plots (Left). The histograms (10° bins) display the percentage of cilia at various angles (Right). **(D)** Changes in cilia length and distribution from P9 to P14. Cyan arrows indicate the direction of cilia re-distribution. Scale bars: 20 µm. **(E)** The percentage of SO-oriented cilia from P10 to P40. **(F)** The thickness of SP lamina at different ages. p < 0.001, one-way ANOVA with Bonferroni’s post hoc test for (E-F). **(G)** Confocal images of calbindin and AC3 at P40. The white boxes are enlarged at the bottom. Scale bars: 10 µm. P: postnatal. Super: superficial sublayer. Deep: deep sublayer. Sub: Subiculum. V3: the third ventricle. n = 3 mice each group.

To test if primary cilia with opposite orientations come from early- and late-born pyramidal neurons, respectively, we performed immunofluorescence staining using a calbindin antibody, a marker for late-born neurons (Soltesz and Losonczy, 2018; Valero et al., 2015). Cilia of calbindin-positive neurons in the superficial sublayer largely orient towards SR, whereas calbindin-negative neurons’ cilia generally were SO-oriented (Fig. 1G). These data indicate that cilia of early- and late-born pyramidal neurons exhibit opposite directions.

### ARL13B Strongly Regulates Neuronal Ciliation in the Hippocampus

ARL13B is an atypical ciliary small GTPase that modulates ciliogenesis and interneuron placement (Gigante et al., 2020; Larkins et al., 2011). Cilia in both WT and ARL13B over-expression transgenic mice were the longest in the CA1 (Fig. S1B), compared to many other brain regions including CA3, dentate gyrus (DG), neocortex, and amygdala (Fig. S1B). We further observed that the expression of AC3 in pyramidal neurons in CA1 SP remained relatively stable after P10 (Fig. 1A). In contrast, ARL13B expression started to emerge at P5 (Fig. 1A). At P10, ARL13B expression reached a peak level, decreasing at P14 and further fading away at P40 and adulthood (Fig. 1A). ARL13B high expression appeared to concur with pyramidal cell positioning from P3 to P14, suggesting a role of ARL13B in positioning.

To examine if ARL13B affects ciliation pattern in CA1 SP, we characterized changes in cilia morphology and orientation during postnatal development using Arl13b+ mice and controls. First, overexpression of ARL13B not only elongated primary cilia, but also affected cilia emanation (Fig. 2A-B). Arl13b+ mice had staining-detectable cilia in the deep sublayer at P3, which were SO-oriented (Fig. S1C). At P7, cilia of deep-sublayer neurons of Arl13b+ mice were very long, while cilia of superficial-sublayer neurons did not fully emanate, with many dot-shaped at SP bottom (Fig. 2A and Fig. S1D). Likewise, cilia in the deep sublayer of Arl13b+ were SO-oriented, whereas those in the superficial-sublayer clearly had opposite direction (Fig. 2A-B). Second, data collected from P7 to P75 indicate that primary cilia in CA1 SP underwent drastic changes in orientation from P7 to P14 and mild changes from P14 to P75 (Fig. 2C-E). These data confirm that the time-window before P14 is crucial for changes in cilia positioning and orientation. Third, Arl13b slightly affected the percentage of cilia orientation: at P14 and P40, more cilia angled toward SO in Arl13b+ mice than in WT mice (Fig. 2D-F). Lastly, immunostaining using calbindin antibody also verified that cilia with opposite orientation were from early- and late-born neurons, respectively (Fig. 2G). These data demonstrate significant changes in cilia positioning and orientation during early postnatal development.

**Fig. 2.**
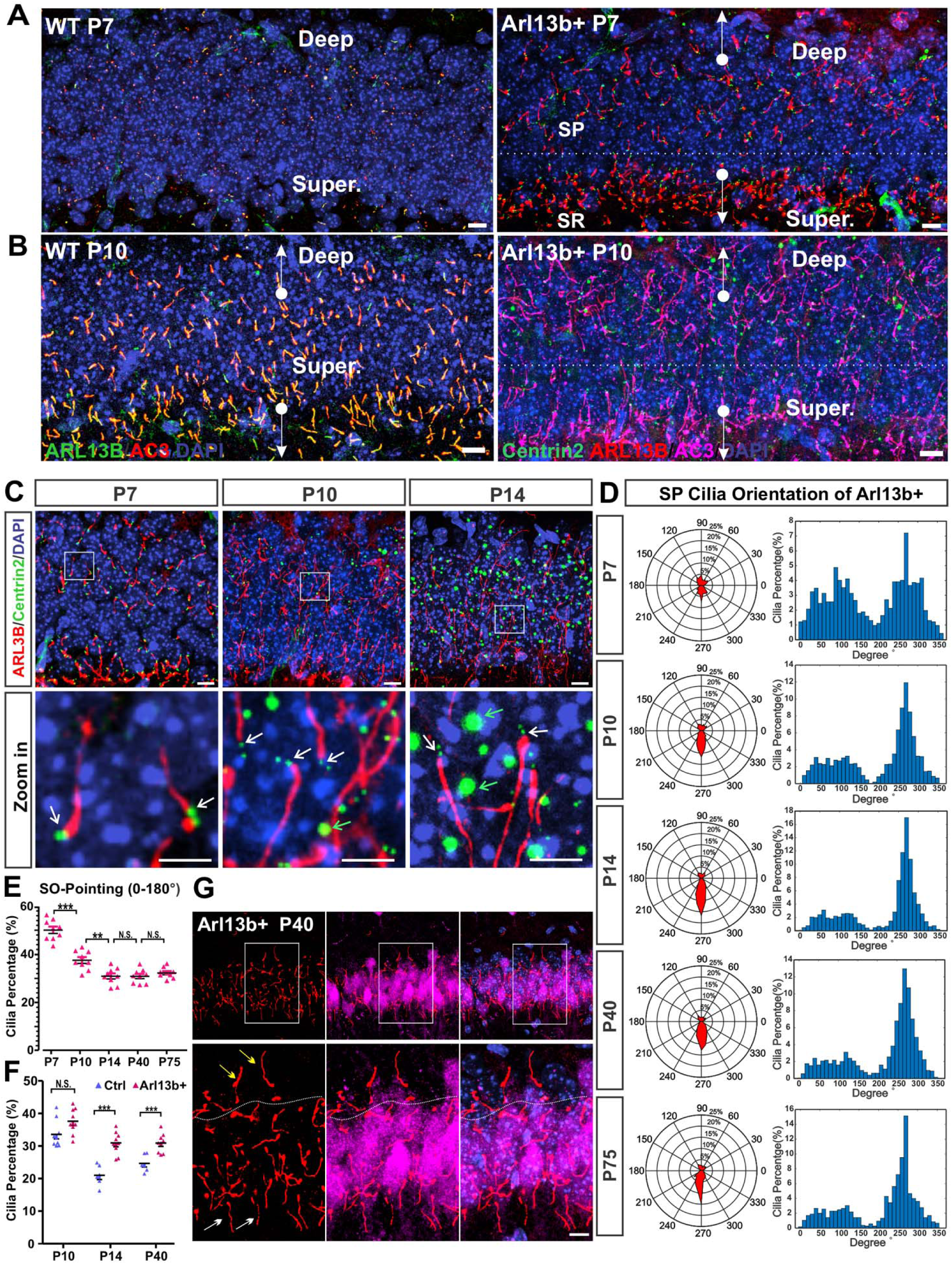
Opposite cilia orientation in CA1 SP of Arl13b+ mice. **(A-B)** AC3- and ARL13B-labeled cilia expression in SP at P7 and P10 in WT (left) and Arl13b+ mice (right) are different. **(C)** Cilia orientation changes at different ages. White arrows indicate the base of cilium, green arrows denote some artifacts. Scale bars: 10 µm top, 5 µm bottom. **(D-E)** The percentage of SO-oriented cilia of Arl13b+ mice decreases drastically during the early postnatal stage. p < 0.0001 by one-way ANOVA with Bonferroni’s multiple comparison test. **(F)** Comparison of the percentage of SO-oriented cilia between WT and Arl13b+ mice. **(G)** Primary cilia of calbindin-positive neurons (purple) at P40 also mostly orient to SR (white arrow), while calbindin-negative neurons’ cilia generally orient toward SO (yellow arrow). Scale bars: 10 µm. White boxes enlarged at the bottom. P: postnatal. Super: superficial sublayer. Deep: deep sublayer. Data were from multiple sections from at least 3 mice each group.

### Neuronal Ciliation Concurs with Neuronal Maturation

Next, we asked the question if neuronal ciliation and maturation occur within the same time-window. We used a NeuN antibody, a marker for matured principal neurons, in an immunofluorescence staining at different ages. We found that the expression of NeuN was barely detected at P1 (Fig. S1E). At P5, NeuN expression became stronger in the deep sublayer, but not in the superficial sublayer of CA1 SP. At P7, the superficial sublayer started to have higher expression, and at P14, NeuN was abundantly expressed in both deep- and superficial-sublayers (Fig. S1E). Next, we searched Allen Brain Atlas Developing Mouse Brain for more evidence. Indeed, many markers of glutamatergic synapses, including Shank3 (Fig. S1F), PSD-95, Synapsin-1 and Syntaxin 1 (Table 1), displayed a similar pattern: first expressed in the deep sublayer of SP at P4, then extended to the superficial sublayer and reached full expression at P14 (Fig. S1 and Table 1). This maturation pattern aligns with the order of cilia emanation from P5 to P14 first in the deep and then superficial sublayer in both WT and Arl13b+ mice (Fig. 1-2).

**Table 1.** Features of neuronal primary cilia and neuronal maturation in hippocampal subregions and the neocortex. Data were collected from multiple sections out of at least 3 mice.

| Regions | CA1<br>-deep | CA1<br>-super. | Cortex<br>-deep | Cortex<br>-super. | CA3<br>-deep | CA3<br>-super. | DG-out<sup>¶</sup> | DG-in<sup>¶</sup> |
| --- | --- | --- | --- | --- | --- | --- | --- | --- |
| Ciliation<br>Time (WT) | P3-P7 | P9-P10 | P3-P7 | P7-P10 | P3-P7 | P3-P7 | P3-P7 | After P7 |
| Ciliation<br>Time<br>(Arl13b+) | P0-P7 | P7-P10 | P0-P7 | P7-P10 | P0-P7 | P0-P7 | P0-P7 | After P7 |
| Cilia<br>Length<br>(WT) | <1 μm,<br>P7 | <1 μm,<br>P7 | 3.3±0.1 μm,<br>P7 | 1.0±0.0 μm,<br>P7 | 5.6±0.1 μm, Adult |  | 2.3±0.0 μm, Adult |  |
|  | 7.0±0.1 μm,<br>Adult |  | 5.6±0.1μm,<br>Adult | 5.0±0.0 μm,<br>Adult |  |  |  |  |
| Cilia<br>Length<br>(Arl13b+) | 1.5±0.0<br>μm, P3 | 1.0±0.0<br>μm, P3 | 1.9±0.1 μm<br>P3 | 0.7±0.0 μm<br>P3 | 8.2±0.1 μm, Adult |  | 3.1 ± 0.0 μm, Adult |  |
|  | 4.5±0.1<br>μm, P7 | 2.8±0.1<br>μm, P7 | 4.5±0.1 μm<br>P7 | 2.1±0.1 μm<br>P7 |  |  |  |  |
|  | 11.7±0.1 μm, Adult |  | 6.3±0.1 μm,<br>Adult | 5.3±0.0 μm,<br>Adult |  |  |  |  |
| Reverse<br>movement | No | Yes | Yes | Yes | TBD |  |  |  |
| NeuN<sup>#</sup><br>Expression | P3-P5 | P7-P14 | P0-P1 | P5 | P3-P5 | P3-P5<sup>‡</sup> | P5-P7 | P7-P10 |
| Camk2a<sup>*</sup><br>Expression | ~P4 | P4-P14 | ~P4 | P4-P14 | ~E18.5-P4 | ~E18.5-P4 | P4 | After P4 |
| Shank3<sup>*</sup><br>Expression | ~P4 |  | ~P4 |  | ~E18.5-P4 | TBD |  |  |
| Homer 1<sup>*</sup><br>Expression | ~P4 |  | ~P4 |  | ~P4-P14 | ~P4-P14 |  |  |
| PSD-95<sup>*</sup><br>Expression | ~P4 or<br>earlier |  | ~P4 or earlier |  | ~E18.5-P4 | ~E18.5-P4 |  |  |
| Synapsin 1<sup>*</sup><br>Expression | ~P4 |  | ~P4 |  | ~ E18.5-P4 | ~E18.5-P4 |  |  |
| Syntaxin1<sup>*</sup><br>Expression | ~P4 |  | ~P4 or earlier |  | ~ E18.5-P4 | ~ E18.5-P4 | Not<br>expressed | Not<br>expressed |
| TBD: to be determined. Super., superficial sublayer. P: postnatal; E: embryonic. Adult: 2–4 months old. <sup>¶</sup> Dentate gyrus outer granular layer and inner granular layer. <sup>†</sup> CA3 deep-superficial distinction in ciliation and neuronal maturation is not as obvious as CA1. <sup>‡</sup> Require further verification. <sup>§</sup> CA3 pyramidal neurons mature several days earlier than CA1 neurons. <sup>#</sup> Based on our immunostaining data using anti-NeuN antibody. <sup>††</sup> Outmost superficial layer in the neocortex <sup>*</sup> Based on ISH data from the Allen Brain Atlas Developing Mouse Brain. |  |  |  |  |  |  |  |  |

However, the CA3 case was different. Both WT and Arl13b+ had cilia expression at P5 and P7 almost at the same time in the deep and superficial sublayers of CA3 SP (Fig. S2A-C). Pyramidal neurons in CA3 SP matured simultaneously and several days earlier than CA1 neurons, as evidenced by NeuN expression and strong expression of many glutamatergic synapse markers in the CA3 at P4 (Fig. S2A-C and Table 1). This pattern sharply contrasts with the weak and deep sublayer-restricted expression in the CA1 initially (Fig. S2D). This suggests that full expression of primary cilia in the hippocampal CA1 region concurs with maturation in an order of “first deep - then superficial”, whereas the CA3 does not display such a pattern.

### Late-born CA1 Principal Neurons Undergo a Reverse Movement to Adjust Position Postnatally

We observed that cilia orientation changed from irregular at P9 upon emanation to highly aligned at P14 (Fig. 1D), and cilia were initially clustered at SP bottom, then become more evenly distributed and well aligned at P14 (Fig. 1D & 2C). We hypothesized that cilia/centrioles reversely move from SP bottom to the main SP. To test it, we examined centrioles/cilia distribution from P3 to P14 using Arl13b-mCherry; Centrin2-GFP mice double transgenic mice (Arl13b+), which mark primary cilia and centrioles, respectively, and Centrin2-GFP single transgenic mice as controls. The density of centrioles in CA1 SP of both Arl13b+ and controls at P3-P7 was quite low (Fig. 3A), and a major portion of centrioles was clustered at the SP bottom (Fig. 3A). We separated CA1 SP into top, middle and bottom panels (Fig. 3E) and quantified nucleus density, centriole density, and the ratio of centriole to nucleus (C/N Ratio) in CA1 SP from P3 to P14 (Fig. 3B-D). First, Arl13b+ mice and controls did not show significant differences in nucleus density at the middle and bottom panels from P3 to P14 in the dorsal CA1 SP (Fig. 3B). However, centrioles were highly condensed at SP bottom at P3 and P7 (Fig. 3A-C), and the C/N Ratio in SP bottom and SR were much higher than 1 (Fig. 3D). However, subsequently at P10 and P14, centrioles showed a relatively even distribution in main SP (Fig. 3C). Consistently, C/N Ratio at P3 was about 2.5, but it decreased to around 1 at P14 (Fig. 3D). We also observed that overexpression of ARL13B in Arl13b+ mice slightly affects the density of cilia clustering in SP, compared to controls (Fig. 3C-D).

**Fig. 3.**
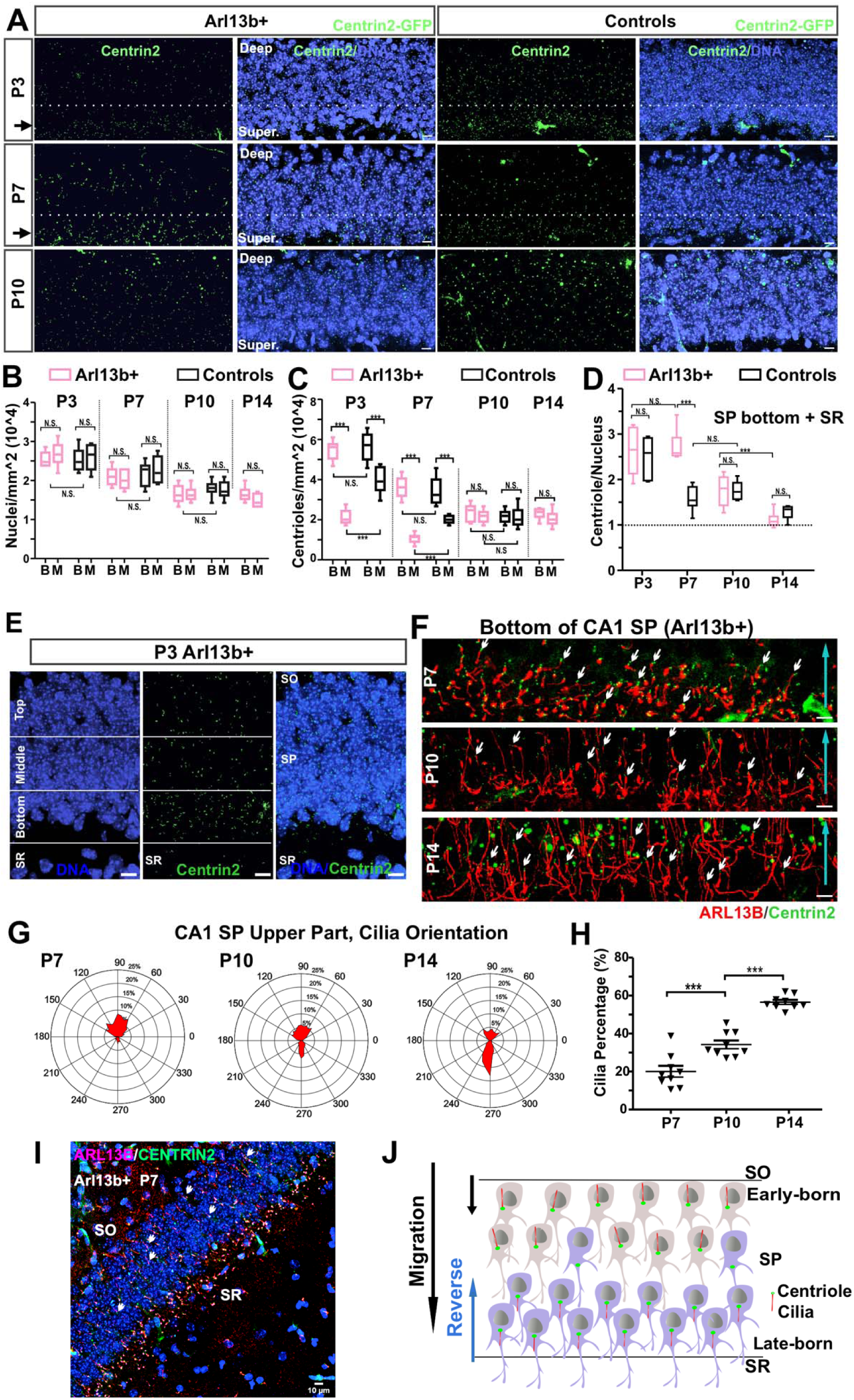
The centrioles/cilia of late-born pyramidal neurons cluster at CA1 SP bottom before reverse. **(A)** The centrioles of Arl13b+ (Arl13b-mCherry, Centrin2-GFP double transgenic mice) and controls (Centrin2-GFP single transgenic) clustered in SP bottom at P3 and P7 but not at P10. Scale bars: 10 µm. **(B-C)** Quantification of the number of nuclei and centrioles (nuclei/mm^2^ and centriole/mm^2^) in the middle and bottom panels. B: bottom; M: middle. Two-way ANOVA with Bonferroni’s multiple comparison test. **(D)** Centriole/Nucleus ratio at SP bottom and SR. **(E)** To quantify centrioles, SP was separated into top, middle, and bottom panels, with a panel of SR of 15 µm. Scale bars: 10 µm. **(F)** Enlarged centrioles/cilia images of late-born neurons showing that cilia length was increasing, and cilia and centrioles were migrating upward (cyan arrows) at P7 and P10. Centrioles in the front were followed by cilia axoneme. Scale bars: 10 µm. **(G)** Polar plots of cilia orientation of upper part of SP at P3, P7 and P10. **(H)** The percentage of SR-oriented cilia in the upper part of SP. p< 0.0001 by one-way ANOVA with Bonferroni’s multiple comparison test. n = 3 mice, 9 sections. **(I)** Arl13b+ centrioles/cilia distribution pattern in the deep and superficial layers of CA1 at P7. **(J)** A diagram depicting principal cell positioning and the reverse movement of centrioles/cilia of late-born neurons.

Moreover, cilia density in the upper part of SP was low at P7, when short cilia were enriched at SP bottom (Fig. 3F). At P10, cilia were elongated. Some centrioles/cilia became well-aligned with the radial axis, while others still clustered at SP bottom (Fig. 3F). Afterwards at P14, cilia occupied the major space of SP, and their cilia more uniformly oriented to the SR direction. A similar pattern was also observed in WT mice (Fig. 1D). We also quantified cilia orientation of the upper part of SP, which only had SO-oriented cilia at P7. More SR-oriented cilia gradually moved up to the main SP from P7 to P14, the upper part of SP subsequently contained more SR-oriented cilia (Fig. 3G-H). Here, the centrioles of late-born principal neurons in the superficial sublayer of CA1 SP move against the direction of radial migration back to the main CA1 SP (Fig. 3I). We name this process “reverse movement”. Given SP lamina thickness reduced correspondingly (Fig. 1F), the change in centriole position and cilia orientation toward SR likely reflects the reverse movement of the soma (Fig. 3I).

### Principal Neurons in the Cerebral Cortex Slightly Move Backwards for Postnatal Positioning

The hippocampal CA1 region and neocortex share a birthtime-dependent inside-out lamination pattern (Soltesz and Losonczy, 2018). To examine whether the neocortex has similarities to CA1 SP, we examined cilia expression, C/N Ratio, and cilia directionality in the postnatal neocortex. Primary cilia were not well protruded in the neocortical superficial sublayer until P7 (Fig. 4A-B). Like CA1 SP, primary cilia were expressed earlier in the deep than superficial sublayer in both WT and Arl13b+ mice (Fig. 4A-D). To determine if neocortical principal neurons are subjected to a reverse movement, we examined centriole/nuclei distribution pattern in the outer layer of the neocortex at P3 and P14 (Fig. 4E-H). We manually drew top and middle panels on the outer layer to quantify centrioles and nuclei density (Fig. 4E). We found that the C/N ratios were near 1 and there were no significant differences between P3 and P14 and between the top and middle panels (Fig. 4F). However, the density of nuclei (Fig. 4G) and centrioles (Fig. 4H) at the top panel were higher than at the middle panel at P3. But at P14, both centriole and nuclei density of two panels had no significant differences. These data indicate that the outer layer change from compact and uneven at P3 to sparse and even at P14. Consistently, the distances from the edge of the superficial sublayer to the pia gradually increase during the first two weeks after birth (Fig. 4I). Because primary cilia in the neocortex before P10 are too short to be accurately quantified (Fig. 4A-D), we thereby quantified cilia orientation in the neocortex using WT and Arl13b+ brain sections at P14 and P40, when primary cilia exhibit a clear directionality. Cilia in both the superficial and deep sublayers oriented toward the pia in WT and Arl13b+ mice (Fig. 4J-K).

**Fig. 4.**
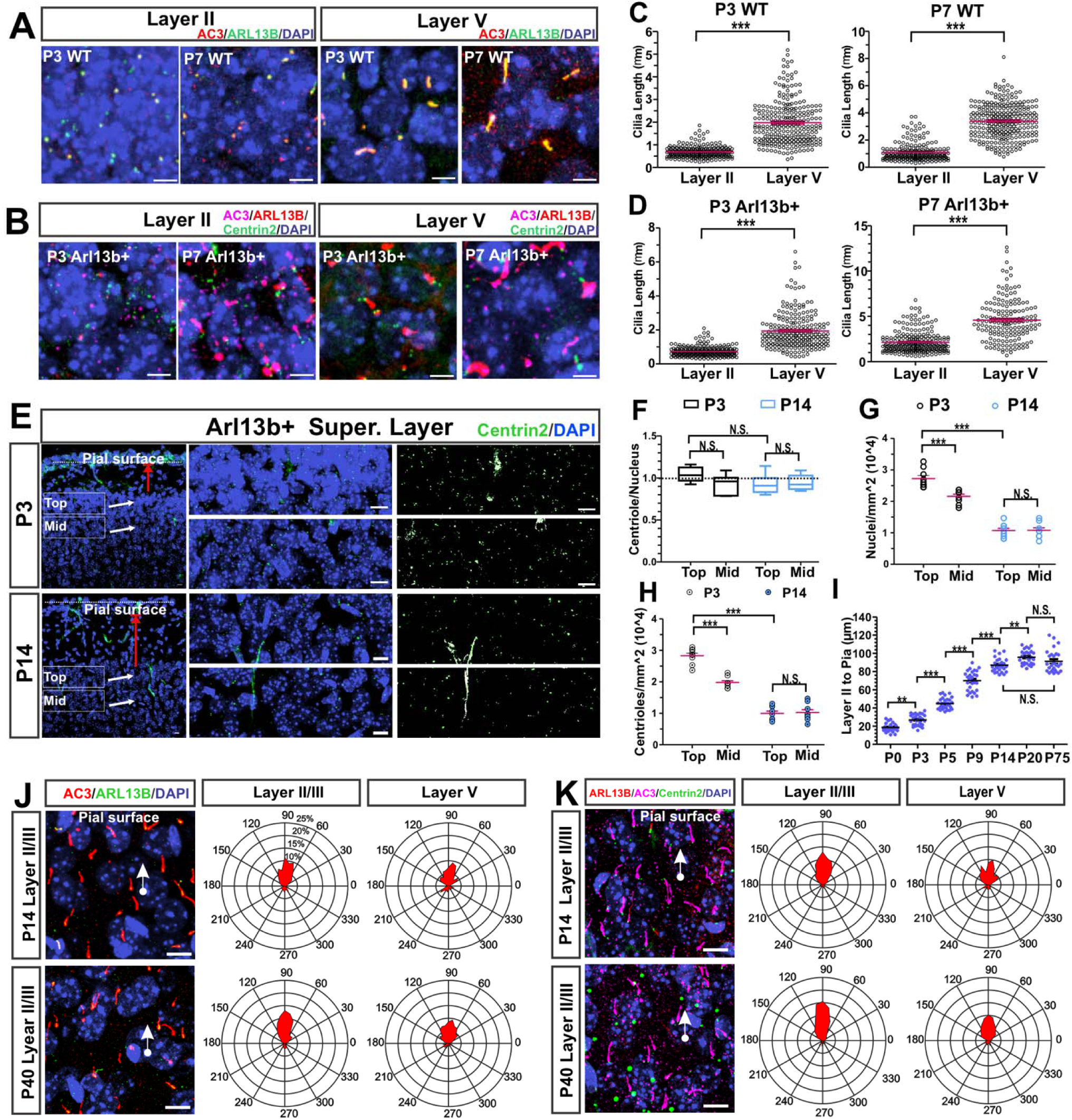
Neocortex cilia orientation suggests a reverse movement of principal neurons during postnatal development. **(A-B)** Co-immunostaining of AC3 and ARL13B in Layer II and V of WT mice **(A)** and Arl13b+ mice **(B)**. Scale bars: 5 µm. (**C-D)** Quantification of cilia length in cortical Layer II and V for **(A)** and **(B)**. Unpaired Student’s T-test. **(E-H)** The densities of centrioles and nuclei in the Layer II/III of the neocortex changed from compact and uneven at P3 to sparse and even at P14. White boxes in the left images are enlarged on the middle. Centrin2 images are shown on the right, with brightness and contrast adjusted. Scale bars: 10 µm. **(F)** The ratios of centriole to nucleus (C/N ratio) in both top and middle of the cortical superficial sublayer at P3 and P14 were all close to 1. The density of nuclei **(G)** and centrioles **(H)** at P3 and P14 were quantified. Two-way ANOVA test with Bonferroni post hoc tests was used in **(F-H)**. **(I)** The distance from the outer edge of Layer II to the pial surface, which is indicated in **(E)** by red arrows, increased significantly in the first two postnatal weeks. p < 0.001 by one-way ANOVA test with Bonferroni post hoc tests. (**J-K**) Cilia in both the superficial sublayer (Layer II/III) and deep sublayer (Layer V) in the neocortex oriented towards the pia at P14 and P40 in WT **(J)** and Arl13b+ **(K)** mice. Scale bars: 10 µm. Data were collected from at least 3 mice.

We next determined primary cilia directionality in other brain regions, including the subiculum, postsubiculum, cingulate cortex, and entorhinal cortex. We found that projection neurons in compact laminae generally had opposite cilia directionality, while neurons in loosely layered laminae predominantly displayed one cilia orientation (Fig. S3A-B, Table 2). We also compared cilia directionality in nucleated brain regions with laminated cortical regions. Cilia in laminated cortical regions, such as the neocortex, hippocampal CA1, subiculum, and entorhinal cortex exhibited a clear directionality: they either had opposite cilia orientations (CA1) or predominantly orient toward the pia (Fig. S3B). However, no specific cilia orientation was identified in many nucleated brain regions, including the thalamus, amygdala, and hypothalamus (Fig. S3A-B, Table 2). We further examined the cilia directionality of interneurons and astrocytes in the hippocampus and neocortex. Primary cilia of interneurons and astrocytes displayed irregular orientations (Fig. S3C-G, Table 2). These data demonstrate that principal neurons in laminated cortical regions display predominant cilia orientations. Additionally, the sharpest cilia directionality manifests at P14 - P20 (Fig. 1-2 and Fig. S3), when principal neurons just complete positioning. Together, these results support hypothesis that the reverse movement of the somata of principal neurons during early postnatal neurodevelopment leads to specific cilia orientation.

**Table 2.** Cilia directionality in different brain regions or cell types, in association with cell density and lamination. Cilia orientation of principal neurons may change to some degree with age. The best timing to measure cilia directionality of principal neurons is at P14-P20 when principal neurons have just completed cell positioning.

| Brain Regions or Cell Types | Primary Cilia Directionality | Cell Density in Lamina |
| --- | --- | --- |
| DG | Opposite | Compact |
| Dorsal CA1 | Opposite | Compact |
| Ventral CA1 | Opposite | Compact |
| Neocortex | Same, oriented to the pia | Loose |
| Cingulate Cortex | Same, to the pia | Loose |
| Entorhinal Cortex | Same, to the pia | Loose |
| Subiculum | Same, to the pia | Loose |
| Postsubiculum | Same, to the pia | Loose |
| CA1 Interneurons | Irregular | Not layered |
| CA1 Astrocytes | Irregular | Not layered |
| Neocortex Astrocytes | Irregular | Not layered |
| Thalamus | Irregular | Not layered |
| Hypothalamus | Irregular | Not layered |
| Amygdala | Irregular | Not layered |
| Striatum | Irregular | Not layered |

To further examine if principal neurons in the neocortex undergo a slow reverse movement during early postnatal development, we quantified the ratio of individual layer thickness relative to the whole six layers as well as the distance from Layer VI to the ventricular zone (VZ). The thickness of superficial sublayer and deep sublayer changed drastically from E18.5 to P20. The superficial sublayer greatly expanded, while the relative thickness of deep sublayer gradually decreased (Fig. S4A-D). In the meantime, the distance from the bottom edge of Layer VI to VZ significantly decreased with age postnatally (Fig. S4E-F). These data support the conclusion that neocortical principal neurons have the space to move backwards.

To further verify that neocortical principal neurons undergo a reverse movement during postnatal development, we utilized CFSE, a cell tracer (Govindan et al., 2018; Telley et al., 2016a; Yoshinaga et al., 2021) to label cells in the outer layer (Layer II/III) of the neocortex at P0 and trace the location of labeled cells several days later. First, CFSE mostly stayed near to injection sites at P1, while many CFSE-labeled superficial neurons appeared in deep region at P3, and further inwards at P9 (Fig. 5A-B). Because CFSE can label daughter cells (Telley et al., 2016b; Yoshinaga et al., 2021), we injected EdU into E18.5 pregnant mice to label new-born cells. We found that CFSE-labeled migrating cells were not merged with EdU (Fig. 5B), excluding the possibility that the CFSE-positive cells were daughter cells of original labeled neurons. Second, to identify the cell type of reversely migrating cells, we used a Ctip2 antibody, an early-born principal neuron marker, to stain CFSE-labeled brain sections. To our surprise, ∼70% CFSE-labeled cells overlapped with Ctip2-marked cells (Fig. 5C-D), suggesting that these CFSE-labeled neurons were excitatory neurons. To rule out the possibility of CFSE diffusion from the injection location, we also injected CFSE into inner cortical layers and found that CFSE did not laterally spread to other regions (Fig. 5E). Together, these data further support the hypothesis that some principal neurons in the neocortex undergo a reverse movement during early postnatal development.

**Fig. 5.**
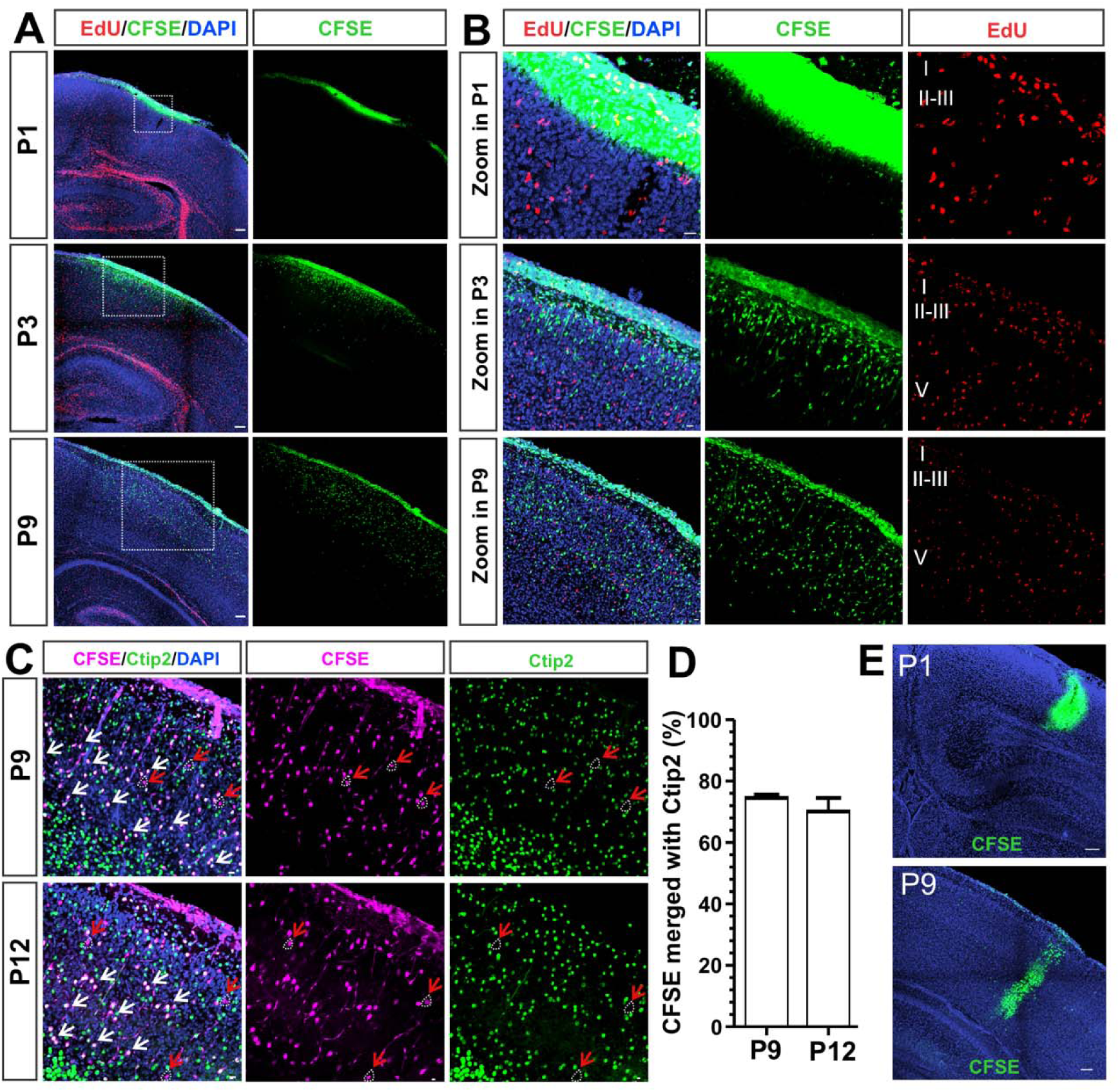
CFSE tracing reveals a slow reverse movement of neocortical principal neurons during early postnatal development. **(A-B)** CFSE tracing shows that principal cells underwent a reverse movement. **(A)** CFSE injected at P0.5, EdU injected at E18.5. CFSE and EdU positive cells were analyzed at P1, P3, and P9. Scale bar: 100 µm. **(B)** White boxes in **(A)** are enlarged. Scale bar: 20 µm. **(C-D)** CFSE-labeled cells were mostly Ctip2-positive cells. White arrows: CFSE merged with Ctip2. Red arrows: CSFE is not merged with Ctip2. Scale bar: 10 µm. **(E)** Control: little lateral diffusion was observed at P1 and P9 when CFSE was injected into neocortical inner layers at P0.5. Scale bars: 100 µm.

We also examined if subiculum and postsubiculum exhibit a reverse movement during early postnatal development by examining Allen Brain Atlas Developing Mouse Brain. We found that the demarcation between the distal CA1 and subiculum was not distinguishable at E18.5 and P4, but the compact CA1 - sparse subiculum distinction became prominent at P14 (Fig. S4G-H). This suggests that subiculum neurons undergo a more drastic reverse movement than CA1 neurons. Moreover, the inward curvature in the postsubiculum was not clear at P4 but became obvious at P14 (Fig. S4, bule arrows). Based on data on cilia/centriole distribution (Fig. 1-3), CFSE tracing (Fig. 5), postnatal changes in cortical lamination (Fig. S4), as well as ISH data from Allen Brain Atlas, we estimated the speed of the reverse movement. While there are strong regional variations, the reverse speed was roughly 0.1-3 µm/hour, which is 10-30 folds slower than typical radial migration (Tabata and Nakajima, 2003). Nevertheless, over time, the slow reverse movement can substantially re-shape laminar structure throughout the cerebral cortex (Fig. S4).

### Alteration of Cilia Function Selective in the Forebrain Affects Principal Cell Positioning and Lamination

Next, we asked if cilia regulate postnatal principal cell positioning. To selectively ablate or shorten primary cilia in the forebrain, we crossed Ift88 floxed strain (Haycraft et al., 2007) and Arl13b floxed strain (Su et al., 2012) with Emx1-Cre mice (Gorski et al., 2002), respectively, which expresses Cre selectively in the forebrain after E12.5. Immunostaining results confirmed that primary cilia in Ift88 flox/flox Emx1-Cre conditional knockout mice (Ift88 cKOs) were deleted in the cerebral cortex but not in other brain regions or non-neuronal organs (Fig. S2F-K), so did Arl13b cKO. We also included Arl13b+ mice, which have elongated primary cilia (Fig. S1B), for a gain-of-function study.

To globally investigate gross anatomical changes, we used X-ray micro-CT to scan the whole brains of P20 Ift88 cKOs and littermate controls. Surprisingly, Ift88 cKO mice manifested gyrification in RSC (Fig. 6A), but not other brain regions. However, this regional gyrification was only observed in Ift88 cKO mice, not Arl13b cKO or Arl13b+ mice. This phenotype was similar to GPR161 cKO (Shimada et al., 2019) and mutant with constitutively active hedgehog signaling (Wang et al., 2016). Secondly, the neocortex thickness of Ift88 cKO mice was significantly higher than controls at P20 (Fig. 6A-D). We quantified the brain weights of Ift88 cKO mice from P0 to adulthood. Ift88 cKO mice did not show significant larger brain size in the embryonic stage (Fig. 6C), which is consistently with previously published data (Snedeker et al., 2017). But a larger brain started to appear at P5 and strong differences between Ift88 cKOs and WTs manifested at P5-P20 (Fig. 6C-D). Additionally, a low percentage of Ift88 cKO mice had spontaneous seizures after P14, lead to some early death (observations). In contrast, Arl13b cKO mice only had slightly increased brain size at P9 but no significant difference at other ages (Fig. 6E). Conversely, the brain size of Arl13b+ mice before P9 was normal, but they had a slightly smaller brain than controls at P14, P20 and P75 (Fig. 6F). These data suggests that cilia signaling modulate postnatal neurodevelopment.

**Fig. 6.**
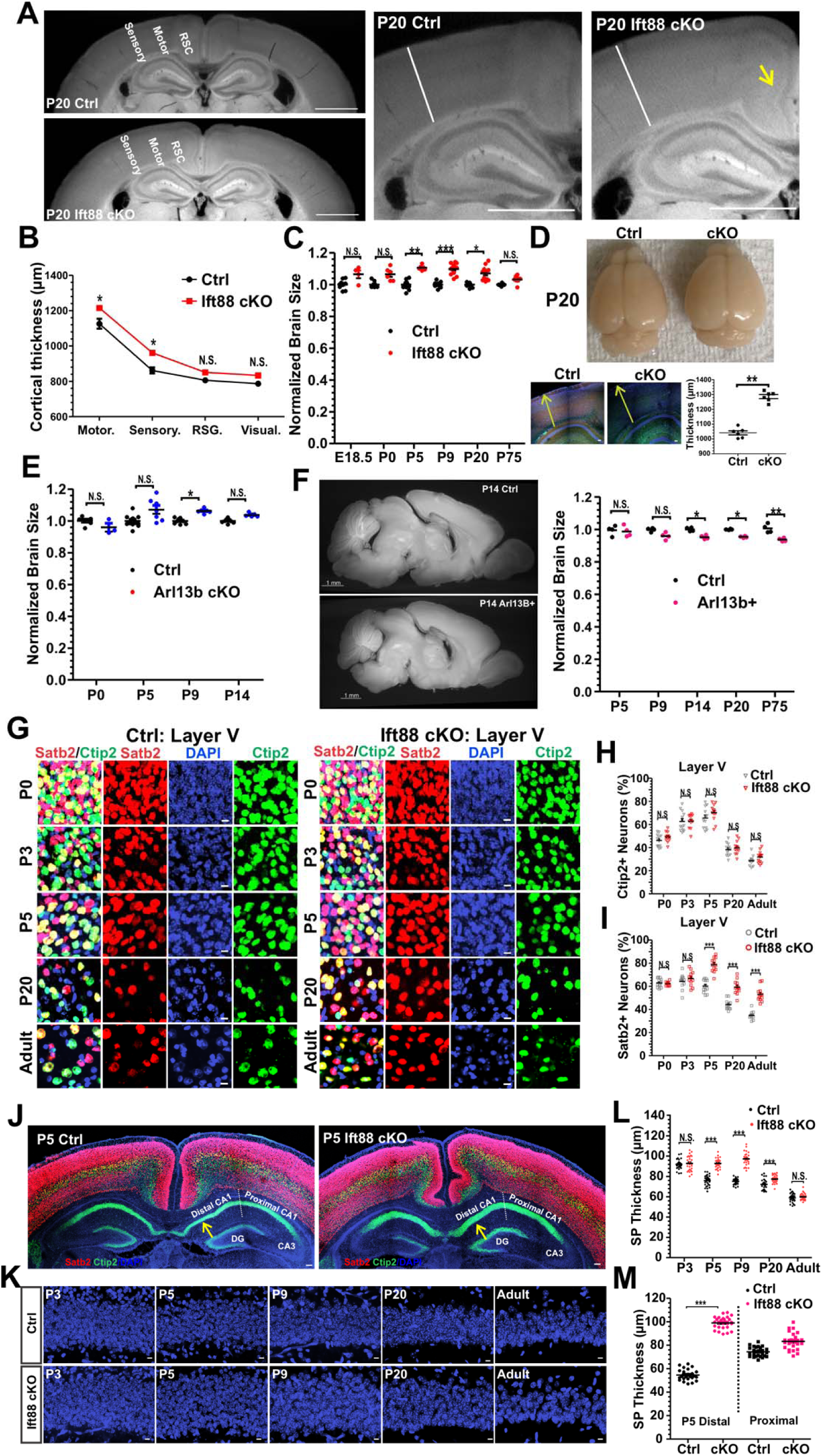
Cilia alterations affect brain volume and lamination. **(A)** X-ray micro-CT images at P20 in control and Ift88 cKO mice. Scale bars: 1 mm. A yellow arrow indicates a gyrification site. **(B)** The thickness of different cortical subregions in control and Ift88 cKO mice was quantified. **(C)** After P5, Ift88 cKO mice displayed significantly larger brain size than controls. One-way ANOVA with Bonferroni post hoc tests. **(D)** Representative pictures of control and Ift88 cKO brains at P20. Lamina thickness was measured in cortical sections in the same location. Two tailed unpaired Students T-test. **(E)** Arl13b cKOs had slightly larger brains at P9, but not at other ages. **(F)** Arl13b+ mice exhibited significantly smaller brain size after P14 than controls (n = 4). Two-way ANOVA with Bonferroni post hoc tests. (**G-I**) Altered neuronal composition in neocortical layers in Ift88 cKO mice. **(G)** Co-immunostaining of Satb2 and Ctip2 antibodies in Layer V at different postnatal ages. Scale bar: 10 µm. (**H**) Ift88 cKO mice had more Satb2-positive neurons (relative to nucleus counts) in Layer V from P5 to adulthood, but not at P0 and P3. (**I**) There were no significant differences in Ctip2-positive neurons in Layer V between Ift88 cKO and control mice from P0 to adulthood. (**J-M**) Cilia ablation alters CA1 lamination. (**J**) Ift88 cKO mice showed difference in the distal CA1. Scale bar: 100 µm. (**K**) CA1 SP thickness decreased with age in both controls and Ift88 cKO mice. Scale bar: 10 µm. (**L**) Quantification of CA1 thickness in control and Ift88 cKO mice. n = 3 pairs for each age group. Two-way ANOVA with Bonferroni post hoc tests. **(M)** The thickness changes of the distal CA1 SP in Ift88 cKO mice were much more severe than those in the proximal CA1 at P5. n = 3.

We also examined neuronal composition in neocortical layers using Satb2 and Ctip2 antibodies, which are markers for late- and early-born principal neurons, respectively. Due to cortical expansion, the neocortical outer layer in Ift88 cKOs had more neurons than controls (Fig. 6A-D and 6J). More interesting phenotype was found at Layer V, which is typically occupied by Ctip2-positive neurons (Fig. 6G-I). There was no difference in Ctip2 neuronal number in Layer V between Ift88 cKOs and controls from P0 to adulthood (Fig. 6H). Nevertheless, Layer V of Ift88 cKOs had more Satb2-positive neurons after P5, but not at P0 - P3, than controls (Fig. 6I). This observation suggests that over-produced neurons in Ift88 KOs move backward to the Layer V, thereby increasing the number of Satb2-positive neurons in Layer V after P3. In addition, Ift88 cKO mice had abnormal CA1 SP lamina (Fig. 6J-M). It is known that CA1 SP thickness gradually decreases with age (Fig. 1F), partly due to apoptosis (Pönniö and Conneely, 2004). Ablation of cilia in Ift88 cKO mice caused delayed lamina thickness reduction during P5-P9 (Fig. 6K-L), when primary cilia started to protrude primary cilia. Notably, the distal CA1 lamina thickness of Ift88 cKO mice had been affected more strongly than that of the proximal CA1 (Fig. 6K&M).

To understand how gyrification in Ift88 cKO mice is formed during postnatal development, we examined the developmental time-course of gyrification at different ages spanning from E18.5 to adulthood (Fig. 7A-B). We found that a shallow gyrus started to appear in RSC and cingulate cortex of Ift88 cKO at E18.5, and the degree of gyrification increased strongly from P0 to P9 and reached a steady state after P9 (Fig. 7A-B). Interestingly, the sulcus was formed via an invagination of the outer layer. We further discovered that not only the degree of gyrification, but also the distance from the midline to Layer II gradually increased with age (Fig. 7A-B), further indicative of neuron reverse movement. Additionally, we measured the distance from the midline to Layer V (marked by Ctip2). Similarly, Layer V early-born neurons in RSC continuously drifted backwards (Figure. 7C-D), as indicated by the increased distance from Layer V to the midline. These data provide additional evidence demonstrating that neocortical principal neurons move backwards during postnatal development.

**Fig. 7.**
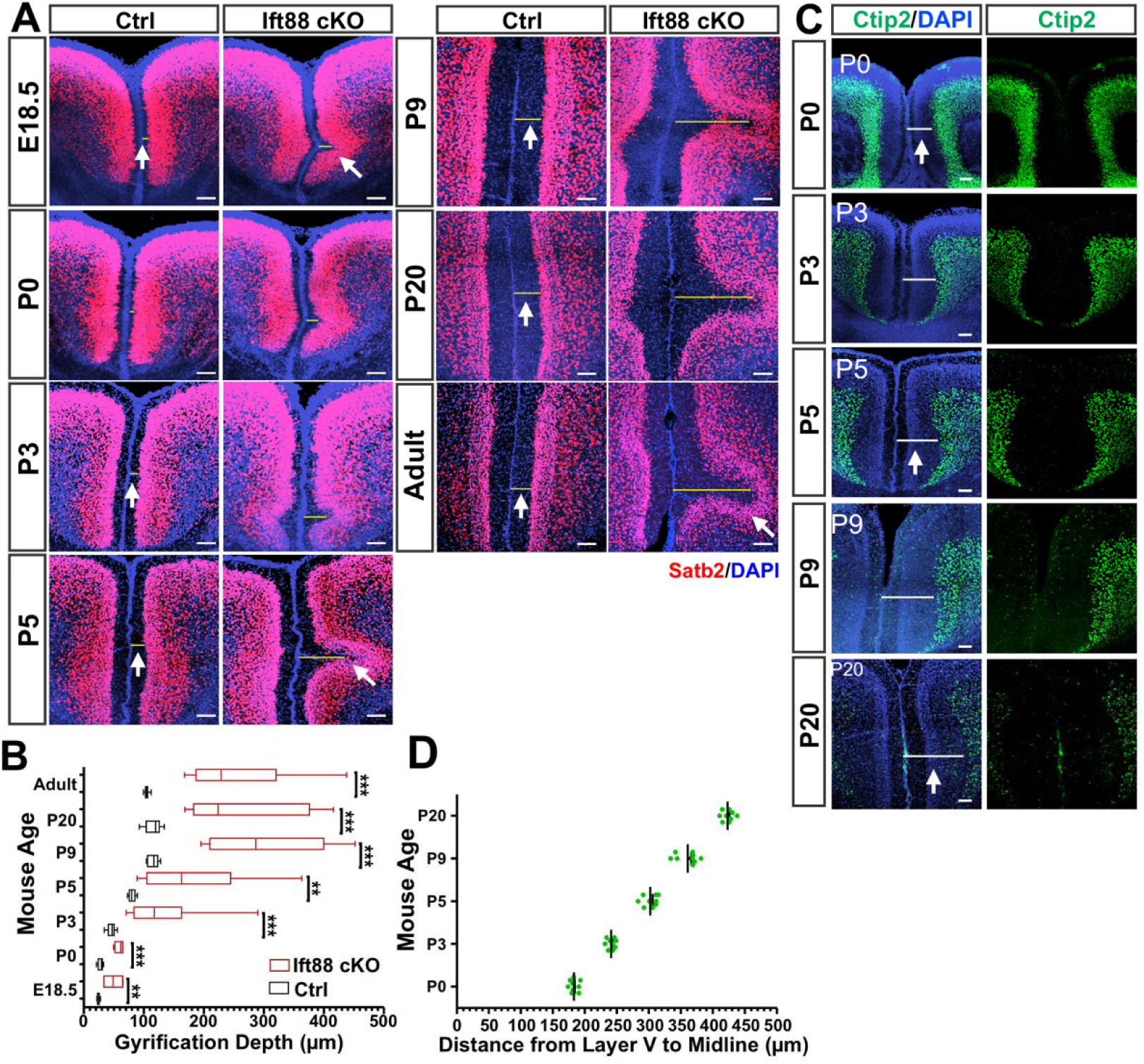
Selective cilia ablation in the forebrain leads to RSC gyrification formed via reverse movement mostly during postnatal development. **(A)** Ift88 cKO mice started to show a very shallow sulcus in RSC at E18.5, and the depth of sulcus increased drastically from P0 to P9. Scale bar: 100 µm. **(B)** The distances from the midline to the edge of the superficial sublayer in controls or to the gyrification site in Ift88 cKO mice significantly increase from E18.5 to P9. Two-way ANOVA test indicated that age affected the distances in both controls and cKOs, p < 0.001 for both ctrl and cKOs, n = 4 - 9 mice, mixed sexes. **(C-D)** The distances from the midline to the edge of the deep sublayer in RSC (marked by Ctip2) from P0 to P20 of WT mice increase. Scale bar: 100 µm. p < 0.0001 by one-way ANOVA test. n = 3 for each group.

Additionally, to evaluate if cilia may modulate neuronal positioning in outside-in laminae, we examined cilia expression pattern and orientation in the granular cell layer of mouse dentate gyrus (DG). Cilia in DG are relatively short (Fig. S1B), thus we leveraged Centrin2-GFP mice, which can mark the base of primary cilia, to help identify cilia orientation in DG. Neuronal primary cilia in the granule cell layer were oppositely oriented at P14 (Fig. 8A-B). Interestingly, there was a developmental shift in ARL13B and AC3 expression in cilia of granular neurons. ARL13B-labeled cilia were prominent at P9 but diminished at P20 (Fig. 8C), when AC3-labled cilia became more predominant (Fig. 8C). Clearly, AC3 is highly expressed in cilia of more matured neurons, while ARL13B tends to appear in cilia of younger neurons (Fig. 8C) (for DG neuronal maturation pattern, refer to Fig. S2D and Table 1). This pattern is reminiscent of the ciliation and neuronal maturation pattern in CA1 SP (Fig. 1) (Fig. S2D), but not in CA3. To test if cilia alterations affect neuronal positioning in the granule cell layer, we analyzed phenotypes of Ift88 cKO and Arl13b cKO mice (Fig. 8D-F). Indeed, cilia ablation (Ift88 cKO) or shortening (Arl13b cKOs) caused neuronal misplacement in the granular cell layer (Fig. 8E-G). The granular cell layer of two strain of cKO mice lost compactness. Many individual Ctip2-positive neurons sparsely distributed in the molecular layer. Of note, the degree of misplaced neurons in the molecular layer in Arl13b cKO mice was lower than in Ift88 cKO mice (Fig. 8G). Together, these data support the notion that primary cilia regulate the postnatal positioning of principal neurons.

**Fig. 8.**
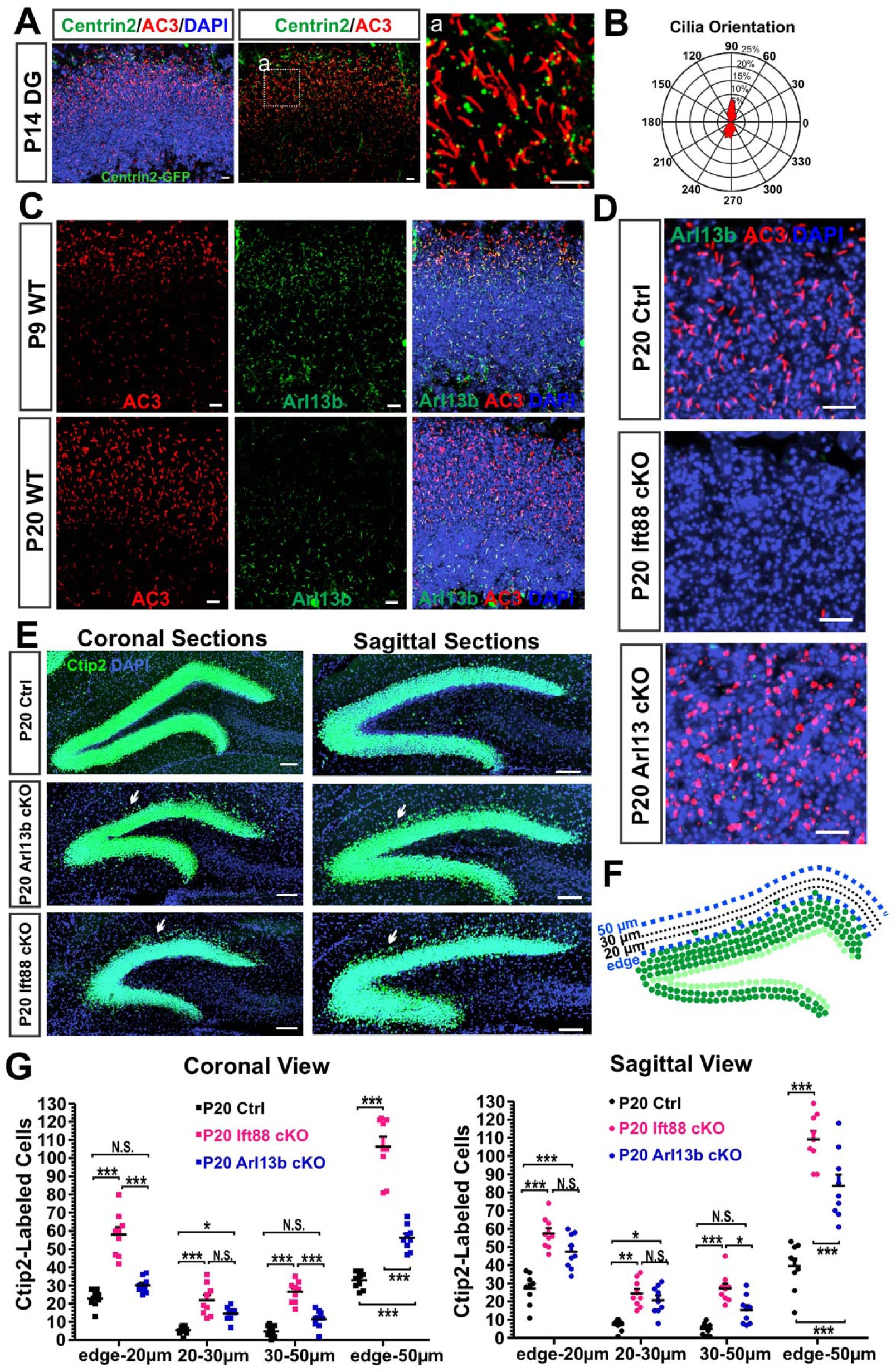
Disruption of cilia function causes neuronal misplacement in DG. (**A-B**) Primary cilia in the granular cell layer are oppositely oriented. Centrioles were marked by transgenic Centrin2-GFP. AC3-labeled primary cilia images in DG at P14 are shown in **(A)** and their orientation was quantified in **(B)**. Scale bar: 10 µm. (**C**) AC3- and ARL13B-labeled cilia at P9 and P20 DG. Scale bar: 10 µm. (**D**) Cilia in DG were ablated in Ift88 cKO mice and shortened in Arl13b cKO mice. Scale bar: 10 µm. (**E**) Ift88 cKO and Arl13b cKO mice show misplaced neurons in DG at P20. White arrows denote misplaced Ctip2-neuorns in DG. Scale bar: 100 µm. (**F**) Quantification diagram of misplaced neurons. (**G**) Statistics of misplaced neurons in DG. Data were collected from 3 mice or 9 brain sections, mixed sexes, two-way ANOVA with Bonferroni post hoc tests.

## Discussion

As the centrosome leads the radial migration during the embryonic development (Higginbotham and Gleeson, 2007; Kuijpers and Hoogenraad, 2011), it is not surprising that the primary cilia of principal neurons do not fully arise until the completion of high-speed radial migration (Matsumoto et al., 2019; Trivedi et al., 2014). After emanation, cilia indicate the positioning of early- and late-born principal neurons to the deep and superficial sublayers of CA1 SP (Fig. 1-3, S4). Cilia expression pattern and cilia directionality support the interpretation: after principal neurons have migrated to the hippocampal plate, primary cilia first protrude out of early-born principal neurons and orient toward SO (Fig. 1-3). With delayed cilia expression, centrioles of late-born neurons migrate a longer distance, pass early-born neurons to reach SP bottom and condense into a cluster. Afterwards, they start to express primary cilia, followed by reverse movement back to the main SP (Fig. 3). It is unclear yet whether the centrioles drive the reverse movement of the soma of principal neurons or simply cilia follow the reversal of soma. Regardless, the movement of late-born neurons is opposite to the direction of radial migration (Fig. 3). The reverse movement refines the position of principal neurons.

It is known that excitatory synapses in the mouse hippocampal CA1 region are labile within the first two weeks after birth and the electrophysiological features of synapses before P14 are different from those after P14 (Fiala et al., 1998). We found that the centrioles of CA1 late-born neurons undergo a substantial reverse movement for positioning (Fig. 1-3). This reverse movement may affect the cytoskeleton of dendrites and the stability of synapses. NeuN staining also revealed that late-born neurons in the hippocampus (Fig. S1D) and the neocortex (Fig. S1G) mature several days later than early-born neurons. These results align well with ISH data of numerous synaptic markers of Allen Brain Atlas. In contrast, primary cilia in the CA3 region does not clearly orient towards specific directions. Neither centriole clustering in superficial sublayer nor marked reverse movement were identified in CA3 SP. Interestingly, CA3 neurons mature earlier and more simultaneously than CA1 neurons (Fig. S2). CA1 and neocortical principal neurons are subject to reverse movement for postnatal positioning during early postnatal development. Therefore, it is tempting to postulate that the delayed maturation of late-born neurons in the CA1 and the neocortex could be, at least in part, due to reverse movement or the delayed completion of cell positioning, while late-born and early-born CA3 principal neurons mature earlier and more simultaneously, partly due to lack of marked reverse movement.

Our analyses of cilia directionality led to the discovery of a slow backward movement of principal neurons during postnatal principal cell positioning. Primary cilia of principal neurons in the cerebral cortex spanning from the neocortex to the CA1 region display specific orientations. Primary cilia oriented oppositely in early- and late-born neurons in the compact CA1 SP as two groups of neurons move opposingly (Fig. 1-2). In contrast, neuronal primary cilia in the neocortical deep and superficial sublayers exhibit a more uniformed orientation (Fig. 4 and S3), suggesting principal neurons reverse move toward the same direction. We have presented multiple lines of evidence showing that principal neurons in the neocortex are subject to a reverse movement for positioning during early postnatal development (Fig. 4-7), so do principal neurons in the subiculum and postsubiculum (Fig. S4). Primary cilia orientations in these areas indicate the direction of the reverse movement (Fig. S3). The mouse brain is composed of both laminated and non-laminated (or nucleated) structures. Laminated structures are typically found in the cerebral cortex. Non-laminated structures are found in other brain regions and do not possess distinct layers. While primary cilia of principal neurons exhibit a directionality in laminated brain regions, they have irregular orientations in non-laminar structures (Fig. S3). In addition, primary cilia of interneurons and astrocytes do not display specific directionality. All converges to support the notion that principal neurons in layered cortical regions display specific cilia directionality, and it is the reverse movement of principal neurons that leads to specific cilia orientations.

One question arises why the cell body of principal neurons undergo a reverse movement for postnatal cell positioning. A simple answer could be that if early-born neurons after arriving the hippocampal or neocortical plates did not reverse to yield some space, late-arrived neurons would locate too close to early-arrived neurons; then all inside-out laminae were only able to form condensed layers, like the CA3 region in most mammalian species (Fig. S5). Following radial migration, the outer layer of neocortex remains highly compacted. Thus far, it is unclear why the mouse neocortical layers are loosely layered, while the hippocampal CA1 and other allocortices become highly condensed. It is interesting that the CA1 SP of many species such as mice and ferrets is highly compact, those of macaques and humans is markedly expanded and loosely layered (Bachevalier, 2019), whereas those of roe deers, jackal and boar are somewhere in between (sparse in the deep sublayer but condensed in the superficial sub-layer) (Malikovic et al., 2022) (Fig. S5). Increased neurogenesis and postnatal neuronal apoptosis may contribute partly to changes in laminar cell density. However, if it were only apoptosis that caused sparse laminae, primary cilia of principal neurons could not always orient toward the pia; the cell density of five principal cell layers in the neocortex would be highly uneven, with very high density in Layer II/III and very low cell density in Layer I-IV. We propose that neuron radial migration stopped by Reelin is not sufficient to construct sparsely layered inside-out laminae spanning from the neocortex to the subiculum in mice or to the CA1 in primates (Bachevalier, 2019) (Fig. S5). An additional step of slow reverse movement, not aways postnatally, is required for proper positioning and lamina refinement.

We propose a working model (Fig. S5), in which a slow reverse movement of principal neurons, coupled with fast radial migration and inside-out lamination pattern, is needed for reshaping lamination in the cerebral cortex. (1) In compact lamina such as the mouse CA1 SP, late-born CA1 principal neurons reverse, whereas early-born neurons generally do not. Early- and late-born CA1 neurons continuously move against each other during postnatal development, gradually decreasing the lamina thickness (Fig. 1F and Fig. S1A). (2) In the subiculum, Ctip2-positive neurons move backwards significantly, encountering little impedance from other neurons. Consequently, subiculum develops into a sparse lamina distinct from the CA1 SP (Fig. S4). (3) In the neocortex, entorhinal cortex, and neighboring regions, both early- and late-born neurons undergo marked reverse movement postnatally (Fig. S4-5), gradually expanding the laminae and forming sparse five principal cell layers (Layer II-VI). Consistently, neuronal primary cilia of early- and late-born neurons orient toward the pia (Fig. 4 and Fig. S3). (4) In some confined cortical regions, such as the RSC or cingulate cortex, over-produced neurons in Ift88 cKO mice are squeezed to undergo a reverse movement to form a sulcus (Fig. 7). (5) We predict that loosely layered CA1 SP of primates (Fig. S5) is likely shaped by a remarkable reverse movement of principal neurons from the condensed superficial sublayer toward the sparse deep sublayer, not necessarily postnatally. (6) In addition to affecting cortical layering, hypothetically reverse movement may impact brain circuit development: the abundance of neurons in the outer layer of neocortex or the superficial sublayer of other cortical regions could serve as a reservoir, from which neurons can gradually move reversely toward deep direction, become matured and form stable synaptic connectivity. This could influence the developmental order of neural circuitry. Together, the reverse movement of principal neurons for positioning can be potentially of great significance in both neurodevelopment and neurophysiology, shaping laminar configuration and modulate synaptic connectivity in the postnatal brain. (7) Although it is best known that the gyrification in the primate neocortex is driven by increased neurogenesis (Fernandez and Borrell, 2023; Ronan and Fletcher, 2015; Ronan et al., 2014), the cellular mechanism of gyrification is not fully understood. Ferret neocortical folding, based on published MRI images of ferret brain development (Barnette et al., 2009), is likely formed by a reverse movement accompanying cortical expansion during postnatal development. We demonstrate here that gyrification is, at least partly, driven by the reverse movement of principal neurons. Together, the concept of reverse movement will open avenues to elucidate how principal neurons complete positioning in the cerebral cortex and provide insights into how 6-layered neocortices evolved from 3-layered allocortices.

## Materials and Methods

### Mice

All experiments are performed using mice with C57BL/6j genetic background. Ift88 floxed mice (Stock No. 022409), Arl13b floxed mice (Stock No. 031077), CaMKIIa-Cre (T29-1) (Tsien et al., 1996), and Emx1-Cre (Stock No. 005628) (Gorski et al., 2002) were purchased from the Jackson Laboratory (JAX). The Arl13b-mCherry Centrin2-GFP double transgenic mice (Stock No. 027967) (Bangs et al., 2015; Higginbotham et al., 2004; Sterpka et al., 2020) were purchased from JAX and originally have C57Bl/6, BALB/c, and FVB/N mixed genetic background. The expression of Arl13b-mCherry and Centrin2-GFP are both driven by the pCAGGs promoter, which contains a chicken β-actin promoter and CMV immediate early enhancer. After importing to the Chen lab, we have backcrossed this strain with C57BL/6j mice for more than six generations to obtain uniform C57BL/6j background. The backcrossing has yielded three separate mouse lines: one keeps as Arl13b-mCherry; Centrin2-GFP double transgenic mice in C57Bl/6 background (Arl13b+ mice), which will be used to mark primary cilia orientation. The other two are Arl13b-mCherry single and Centrin2-GFP single transgenic mice. Centrin2-GFP was used to mark centrioles which locate to the base of primary cilia. Mixed males and female mice were used in the experiments. Mice were maintained on a 12-hour light/dark cycle at 22 °C and had access to food and water ad libitum. All animal procedures were approved by the Institutional Animal Care and Use Committee at the University of New Hampshire (UNH).

### Brain Tissue Collection

Adult mice were deeply anesthetized with an intraperitoneal injection of ketamine and xylazine, and then were transcardially perfused with ice-cold 0.1 M phosphate buffer saline (PBS) followed by 4% paraformaldehyde (PFA) in 0.1 M PBS. Young pups (P0 - P7) were directly decapitated using a sharp surgical scissors. Mouse brains were extracted and then placed into 4% PFA overnight at 4 °C. The following day, brains were rinsed in 0.1 M PBS for 30 min at room temperature. P0-P5 mouse brains then were dehydrated by 30% sucrose in 0.1 M PBS overnight at 4°C. Finally, those tissue were embedded in O.C.T resin before being sectioned at -18 °C using a cryostat to a thickness of 30 μm. P7 - P75 mouse brains were coronal sectioned using Compresstome (Precisionary Instruments LLC, NC) to a thickness of 30 - 50 μm.

### Immunofluorescence Staining

E18.5-P5 mouse brains were sliced and mounted on gelatin-coated slices for immunostaining procedure, while P7 – P75 brain sections were sliced and placed in 24-wells plate, staying afloat during the staining procedure. They were washed three times with 0.1 M PBS. Then they were washed in 0.2% Triton X-100 (vol/vol) in 0.1 M PBS three times for 10 minutes at room temperature. Then they were blocked in blocking buffer (0.1M PBS containing 10% normal donkey serum (vol/vol), 2% bovine serum albumin (weight/vol), 0.2% Triton X-100 (vol/vol)) for one hour at room temperature. After that, slices were incubated with primary antibodies overnight at 4 °C in blocking buffer. After washing in 0.1 M PBS for three times for 10 minutes, slices were incubated with secondary antibodies for 2 hours at room temperature. Primary antibodies were rabbit anti-AC3 (1:10000, EnCor Biotechnology Inc., #RPCA-AC3), mouse anti-ARL13B (1:500, UC Davis clone N295B/66), mouse anti-calbindin (1:1000, Swant Switzerland, D-28K), rat anti-Ctip2 (1:500, Millipore-Sigma, MABE1045), rabbit anti-Satb2 (1:500, Abcam, ab92446), and mouse anti-NeuN (1:500, Millipore-Sigma, MAB377, clone A60). Secondary anti-bodies were Alexa Fluor 488-, 546-, or 647-conjugated (Invitrogen, La Jolla, CA). Finally, sections were counter-stained with DAPI (Southern Biotech, #OB010020) and images were acquired with confocal microscope (Nikon, A1R-HD), installed in the University Instrument Center at UNH.

### Cilia Orientation Measurement

When a single primary cilium in WT mice was measured, the relatively thinner axoneme was defined as the tip and the opposite site as the base. For WT mice, if some cilia were too short to quantify their orientation, or cilia had a uniform axoneme, their orientation were not determined (4.74% for p10, 6.68% for p14). For Arl13b-mCherry Centrin2-GFP transgenic mice, the location of the primary cilium base was the same as Centrin2-GFP. The tails of most primary cilia in WT mice were straight and easy to determine their angles. However, primary cilia in the Arl13b-mCherry Centrin2-GFP transgenic mice were much longer and showed kinks in their axoneme (Sterpka et al., 2020). For those primary cilia, the initial part (∼2 µm from the base) was used for determining the angle of those primary cilia. Fiji was utilized to measure the orientation of the single primary cilium. Four steps were used to complete a measurement. 1) First, the angle tool was selected among icons showed in the main window. 2) Then, the shift button on the keyboard was held, while left clicking the mouse meanwhile to draw a line (always from the right to left and parallel to the external capsule) to the location of the base of the primary cilium. 3) Afterwards, the shift button was released, and the mouse was left clicked to draw a line from the base to the tip of the primary cilium. 4) Lastly, while holding the control button and selecting “M” on the keyboard, the angle would jump out in a separate window. To measure the next primary cilium, steps 2-4 were repeated. Acquired data were finally exported as an excel file and then to a MATLAB program, which was used to generate polar plots.

### Primary Cilia Length Measurement

We measured the length of primary cilia based on published methods (Sipos et al., 2018), with modifications. The freehand line tool within Fiji (ImageJ) was used to measure cilia length from the base of axoneme to the distal tip.

### CA1 SP Thickness Estimation

Confocal images of 20 μm Z-stack were used for quantifying CA1 SP thickness for both WT, Arl13b+ mice and Control mice. The straight-line tool within Fiji was used to measure CA1 SP thickness from the top of SP to the bottom of SP based on nucleus staining (Ponnio and Conneely, 2004). Because the nuclei at the top and bottom edge of SP are not aligned to a straight line, this causes some variation of SP thickness. For young adult mice, we chose coronal sections at -1.9 mm posterior to Bregma to measure CA1 SP thickness. For young pups, we measured the thickness based on selected slices with hippocampal anatomic geometry similar to adult hippocampal sections. We acknowledge that our estimation method has limitations and stereotaxic accuracy needs to be improved. Nevertheless, it is known (Ponnio and Conneely, 2004) that CA1 SP thickness decreases markedly with age, particularly during the first two weeks. We present CA1 SP thickness changes with ages to demonstrate that hippocampal principal cells maintain certain movement or re-positioning postnatally, yielding significant changes during early postnatal period (before P14) and mild adjustment afterwards.

### Quantification of Nucleus Density, Centriole Density, and Centriole to Nucleus Ratio (C/N Ratio)

A 70 µm x 70 µm square box in SP was used for one single area quantification. The freehand line tool within Fiji was utilized to draw a line that matched the margin of the nucleus at the bottom of SP in the chosen box. This line was then vertically moved down 15 µm (to the SR direction) and up 15 µm (to the SO direction). The panel between two lines was defined as the bottom panel of SP for quantifying nucleus, centrioles, and C/N Ratio. The top line at the bottom panel of SP was further moved up 30 µm (to the SO direction). The area between the two lines (having a width of 30 µm) was defined as the middle panel of SP for the quantification of nuclei and centrioles. Since many centrioles were found to be clustered in the SR particularly in Arl13b+ mice, we also included a 15-µm-width area in the SR that was next to SP for quantification. Two Centrin2 dots were counted as one pair of centrioles, one DAPI separate round region were counted as one nucleus.

### Normalized Brain Size/Weight

Due to the variation of maternal care and individual body sizes, the same batch of pups had variations in body weights and brain size. Thus, control littermates, cilia KO mice, or Arl13b+ mice differed by less than 10% in body weight were used to compare their brain size. To normalize brain size at different ages, we first averaged the whole brain weights in controls from the same cage, then divided each mouse brain weight by this average number. Then a final normalized brain size for each mouse was completed.

### Cortical Injection of CellTracer

CFDA-SE is carboxyfluorescein diacetate succinimidyl ester, whose acetyl group is responsible for the initial membrane permeability of the compound and is cleaved by intracellular esterases upon entry into the cell, giving rise to CFSE. CFSE is fluorescent and accumulates intracellularly and can maintain fluorescence for over one month. To label cells in the neocortex of neonatal pups, pups had 0.5 ul of 0.25 mM CFSE dye (Life Technologies, #C34572 and Dojindo Molecular Technologies, #C375) injected into the neocortex, covering the pial surface. The CFSE dye was colored with Fast Green to monitor successful injection. Pups were anesthetized by hypothermia by laying them on an ice bed (icebox filled with ice and cover it with aluminum foil. Anesthesia was assessed by the loss of movement. P0-1 pups would need 1–2 min to be anesthetized by laying it on top of the ice bed, P3 and P5 pups need 3–6 min on the ice bed (Oomoto et al., 2021). The head of the anesthetized pup was placed in a head holder (Olivetti et al., 2020) and secured with a small piece of cloth tape. An injection needle or glass pipette which contained CFSE dye was inserted perpendicular to the cortical layer. CFSE dye was manually injected into the neocortex by slowly pushing the injection bottom on the pipette. If the injection was successful, the colored dye slowly spread under the skull, which were visible. The whole injection procedure were less than 8 minutes. Afterwards, pups were removed from the head holder and placed inside the recovery box on top of a heating mat for 10 minutes to achieve the normal body temperature, before gently returning the injected pups to their mother with the bedding.

### EdU Injection and Staining

EdU (5-ethynyl-2-deoxyuridine, Carbosynth #: NE08701) was dissolved in saline at 1 mg/ml and stored at -20. E18.5 pregnant mice were given one-time intraperitoneal injection of EdU (100 mg/kg). For staining EdU, fixed brain slices were permeabilized for 30 minutes in PBS + 0.2% Triton X-100, and rinsed in PBS for 10 minutes, and then incubated in the fresh cocktail (Tris-buffered saline (100 mM final, pH 7.6), CuSO4 (4 mM final, Acros #197730010), Sulfo-Cyanine 3 Azide (2-5 uM final, Lumiprobe, #D1330), and fresh Sodium Ascorbate (100 mM final, freshly made each use, Acros, #352685000)) for 30-60 minutes. Lastly, rinsed brain slices 3 times in PBS for totally 30 minutes.

### X-ray Micro-CT Scanning of Iodine-Stained Brains

Mice were anesthetized and transcardially perfused with ice-cold PBS, followed by ice-cold 4% PFA in PBS. Brain tissues were carefully dissected and post-fixed in 4% PFA in PBS for 3 days at 4°C. Brains were dehydrated in ethanol grade series (30%, 50%, 70%, 80%, 90%) for 2 h followed by 7 days staining in 90% methanol plus 1% iodine solution at room temperature in a dark area (replace with fresh staining buffer every 2 days.). After staining, brains were rehydrated with 30% and 70% ethanol, for 1 hour each, at room temperature with mild agitation. Finally, brains samples were fixed in 1% agarose gel (Apex Bioresearch Products, #: 20-102GP) and placed in 5 ml tubes at 4°C. All iodine-stained mouse brains were scanned using Zeiss 610 Versa X-Ray micro-CT scanner. The micro-CT scan was carried out in an air-conditioned cabinet at 80 kV acceleration voltage and 125 μA tube current, 0.16° rotation step through 360 degrees, giving rise to 2201 projections; no filter was used. Exposure time was 6 seconds, and 3 X-ray projections were acquired in each angle increment and averaged for reduction of noise. The achieved resolution was around 6 μm with isotropic voxels. Tomographic reconstruction was performed using Dragonfly 2020 software.

### Quantification of Misplaced Ctip2-stained Neurons in DG

We quantified misplaced Ctip2-neurons in the molecular layer of DG as indicated in Fig. 9F. Fiji was utilized to count the cell numbers. Firstly, the freehand line was selected among icons showed in the main window, then drawing a line which almost adjoined and covered the full outmost edge of granule cell layer (GCL) in the top layer and selecting “M” on the keyboard. Next, drawing several lines which is 20 μm, 30 μm, 50 μm perpendicular to the first line. Then copying the first line and move it parallelly to the end point of those 20 μm, 30 μm, 50 μm lines. After finishing this area selection step, the Plugins > Analyze > Cell Counter was selected among icons showed in the main window, after cell counter window showed up, first initializing the image, and then counting.

### Software, Statistical Analysis, Data Presentation, and Manuscript Writing

Data were analyzed with Fiji, GraphPad Prism and MATLAB. Data were considered as statistically significant if p < 0.05 and values in the graph are expressed as mean ± standard error of the mean (SEM). Statistical analyses were conducted using one-way or two-way ANOVA for multiple group comparison or unpaired Student’s T-test with a two-tailed distribution, as appropriate. If not otherwise indicated in the text or Figure legends, N.S., Not Significant; * p < 0.05; ** p < 0.01; *** p < 0.001. **** p < 0.0001. Data were collected from multiple sections from 3 or more mice (mix sex) each group, if not otherwise indicated. Primary cilia directionality analysis and knockout mouse phenotyping studies have been separately repeated by three graduate students (Juan Yang - first author, Soheila Mirhosseiniardakani - second author, Sumaya Akter- not a co-author in this manuscript but performing another project on neuronal positioning). Three undergraduate students (Kostandina Bicja, Abigail Del Greco, Kevin Jungkai Lin) helped repeat many experiments and conducted data analysis. We wrote the manuscript but used OpenAI ChatGPT to correct grammatic errors or typos.

## Funding and Acknowledgments

This study was supported by National Institutes of Health Grants K01AG054729, P20GM113131-7006, R15MH126317-01, and R15MH125305-01 to X.C.; UNH CoRE PRP awards and Cole Neuroscience and Behavioral Faculty Research Awards to X.C.; UNH Summer TA Research Fellowships (STAF) to J.Y and S.M.; a Dissertation Year Fellowship (DYF) to J.Y.; and a SURF award from the UNH Hamel Center for Undergraduate Research to D.K. We would like to thank Brendon Lewis for helping analyze cilia directionality in the ventral CA1 region. We are grateful to the University Instrumentation Center at UNH for A1R HD confocal imaging service.

## Competing Interests

The authors declare no competing interests.

## Supplemental Information

### Supplemental Figures and Figure Legends

**Fig. S1.**
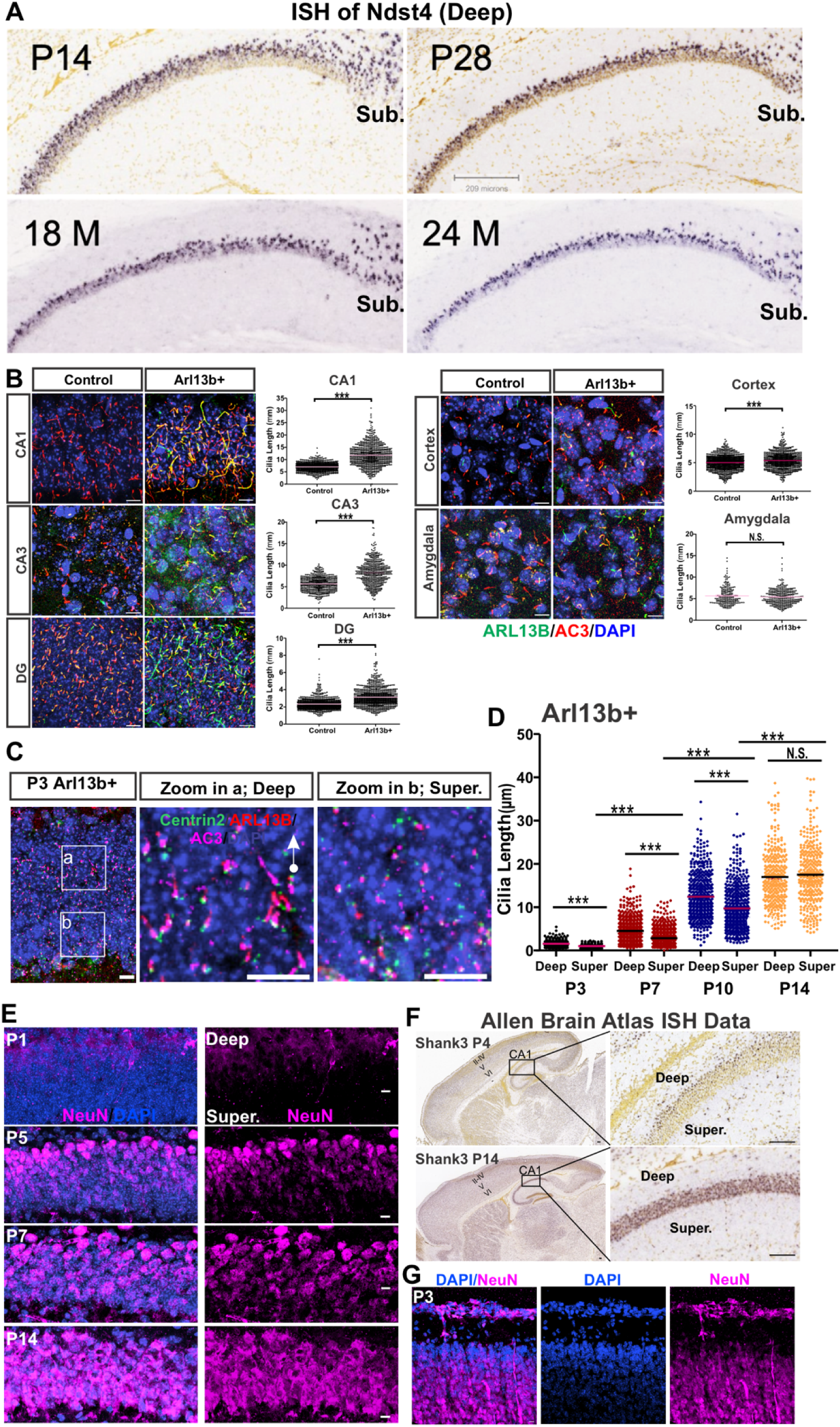
Neuron ciliation coincides with maturation in CA1 SP. **(A)** The thickness of the CA1 SP decreases with age. The distribution of mRNA of Ndst4 (a deep sublayer marker) was dynamic with age. Data were adopted from Allen Brain Atlas. M: month. **(B)** Strong regulation of CA1 neuronal cilia length by ARL13B. Arl13b+ mice had much longer neuronal primary cilia than WT mice in hippocampal regions (CA1 and DG) in adults, but not much in the neocortex and amygdala. Scale bar: 10 µm. Data were collected from at least 3 young adult mice for each genotype, mixed sexes. Two tailed unpaired Student’s T-test. **(C-D)** Primary cilia first emerge from deep-sublayer neurons. **(C)** At P3, early-born neurons in the deep sublayer protruded cilia earlier than neurons in the superficial sublayers of the CA1 in Arl13b+ mice, and those early-protruding cilia oriented towards SO. **(D)** Cilia length of deep- and superficial-sublayer neurons of Arl13B+ mice at P3, P7, P10, and P14. At P14, cilia length reaches the steady-state and has no difference between two sublayers. Two tailed unpaired Student’s T-test; *** p < 0.001, N.S., not significant. **(E)** NeuN expression was first detected in deep-sublayer of SP at P1, more at P5, then extended to superficial-layer neurons at P7. At P14, both deep- and superficial-layer neurons had abundant NeuN expression. Scale bars: 10 µm. **(F)** Allen Brain Atlas in situ hybridization (ISH) results of Shank3, a synaptic maker of excitatory neurons, in hippocampal CA1 at P4 and P14. At P4, Shank3 was only expressed in the deep sublayer of SP and Layer V (deep sublayer) of the neocortex. At P14, ISH signals of this gene were presented in most areas of SP and the six layers in the neocortex. Scale bars: 100 µm. **(G)** NeuN staining in the postnatal neocortex. Principal neurons in outer layer of the neocortex are not matured at P3. In addition, NeuN expression abundance display a decreasing gradient from the deep to superficial layers, even neuron numbers increase.

**Fig. S2.**
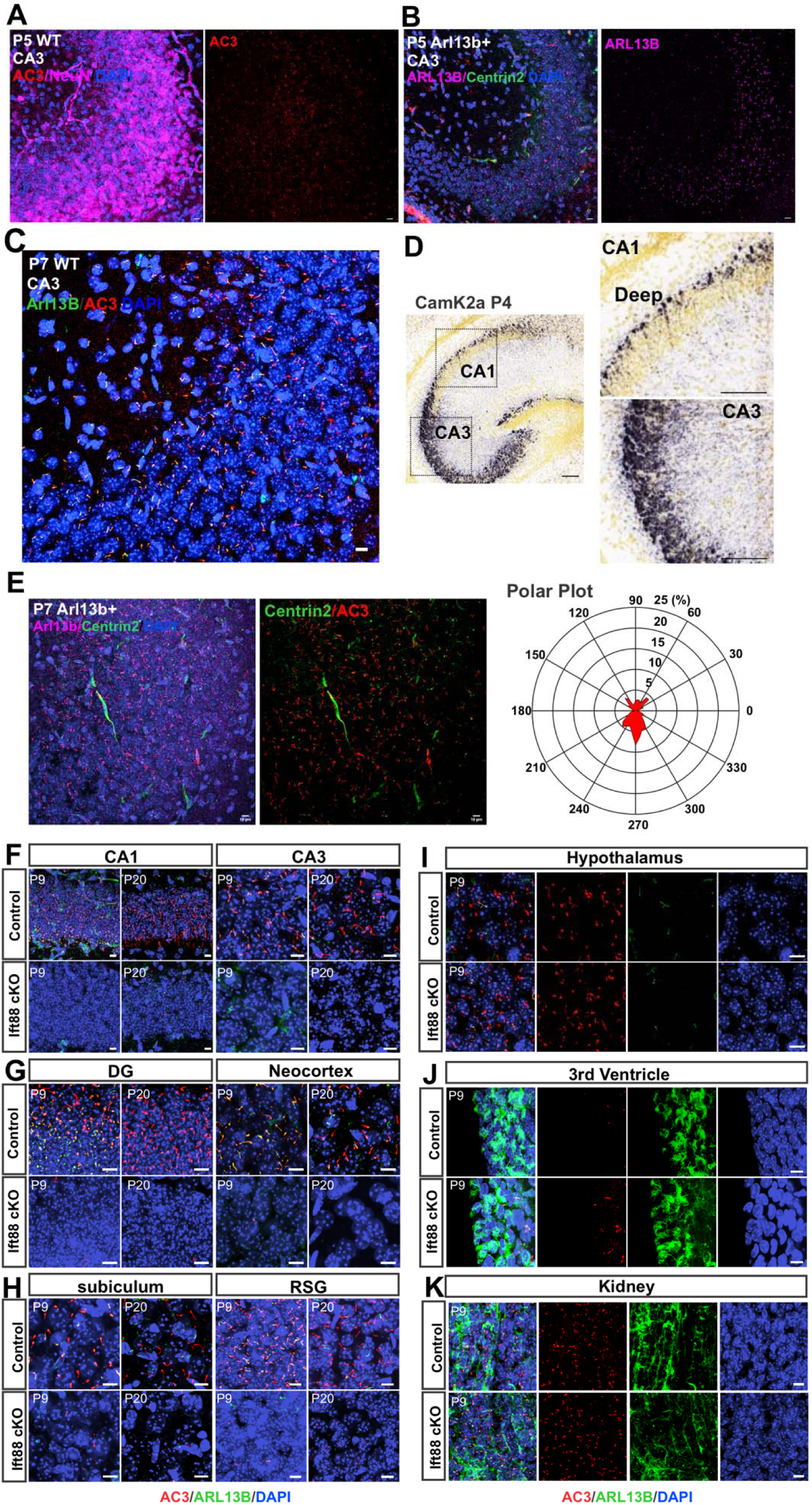
(A-E) CA3 neurons mature and protrude cilia earlier than CA1 neurons. (**A-C**) Neuronal primary cilia were abundantly expressed in the CA3 region in controls and Arl13b+ mice at P5 and P7. NeuN expression in CA3 at P5 **(A)** was strong. Scale bars: 10 µm. (**D**) CA3 pyramidal neurons matured earlier than CA1 neurons. At P4, CamK2a, a postsynaptic marker protein, was highly and widely expressed throughout the CA3. In contrast, its expression was only restricted to the deep sublayer of CA1, not yet present in the superficial sublayer. Scale bars: 100 µm. ISH images were adopted from the Allen Brain Atlas Developing Mouse Brain. (**E**) Cilia directionality in the CA3 region is different from the CA1. Confocal imaging was collected from Arl13b+ mice. **(F-K) Primary cilia are selectively ablated in the forebrain in Ift88 cKO mice.** Primary cilia were ablated in the hippocampal and neocortical regions of Ift88 cKO mice at P9 and P20 (**F-H)**. RSG: granular retrosplenial cortex area, DG: dentate gyrus. Primary cilia were maintained in other regions of Ift88 cKOs including the hypothalamus, 3rd ventricle, and kidneys **(I-K)**. Scale bars: 10 µm.

**Fig. S3.**
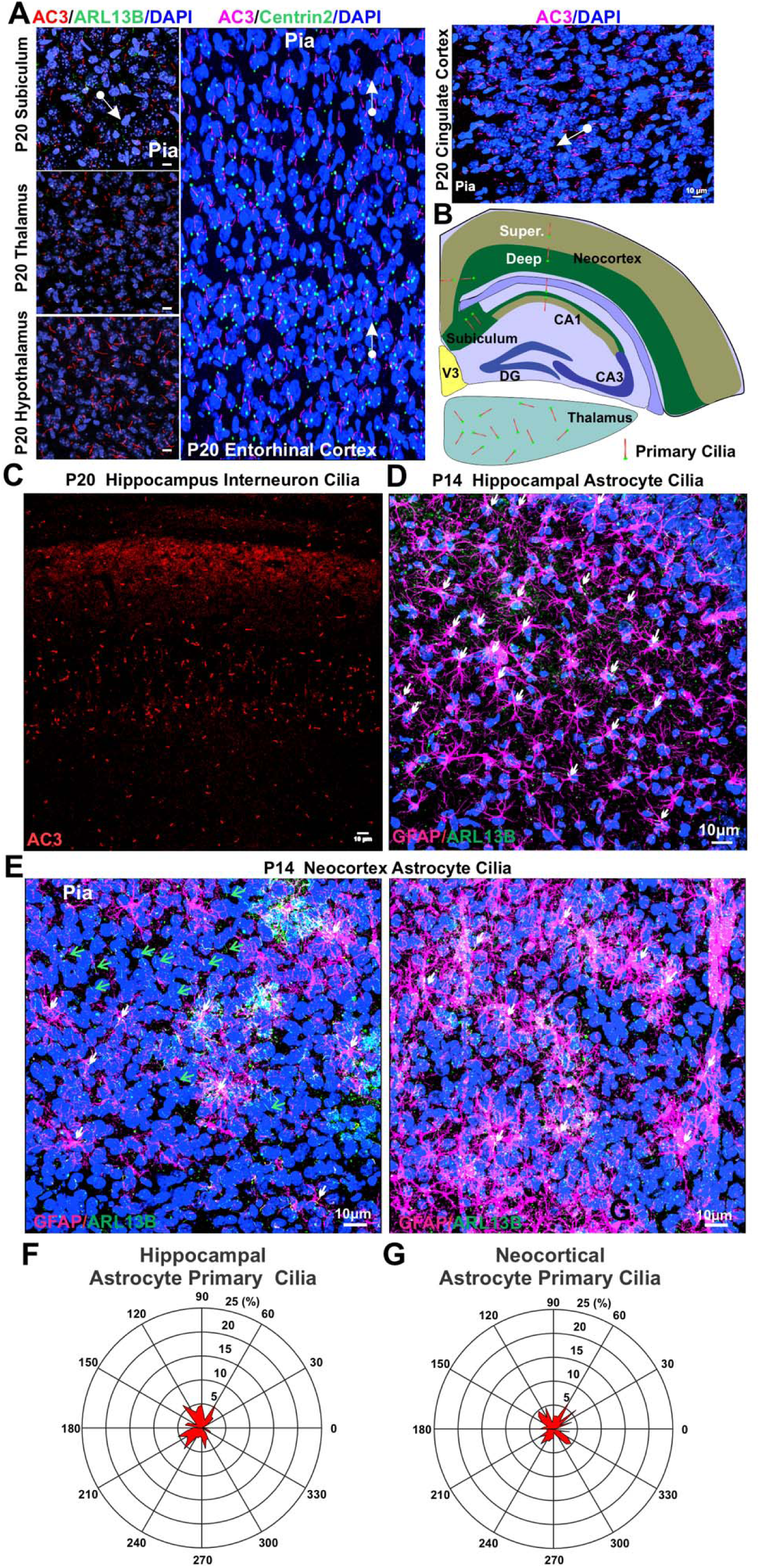
Only principal neurons in laminated cerebral cortical regions exhibit specific cilia directionality. **(A)** Immunostaining of primary cilia in different brain regions including subiculum, thalamus, hypothalamus, entorhinal cortex, and cingulate cortex. Scale bar: 10 µm. **(B)** A diagram summarizing the directionality of neuronal primary cilia in multiple brain regions. **(C-G)** Primary cilia of interneurons and astrocytes do not display specific directionality. **(C)** Interneurons’ primary cilia do not orient to specific directions. Straining using Ift88 flox/flox Camk2a-Cre (T29-1) P20 sections shows remaining primary cilia of interneurons in the hippocampal CA1 and neighboring neocortex. Because AC3 is a marker for neuronal primary cilia and the Camk2a-Cre strain has Cre expression selectively in forebrain principal neurons, primary cilia marked by AC3 in Ift88 flox/flox CaMK2a-Cre belong to interneurons. **(D-G)** Astrocytic primary cilia in the hippocampus (**D**) and in the neocortex (**E**), denoted by white arrows, did not exhibit specific directionality. White arrows denote astrocyte cilia, which display irregular pointing. Green arrows denote neuronal primary cilia, which orient toward the pia. (**F-G**) Polar plots of astrocytic primary cilia orientation in the hippocampus (**F**) and neocortex (**G**). Data were collected from Arl13b-mCherry mouse brain sections that were stained with ARL13B and GFAP antibodies.

**Fig. S4.**
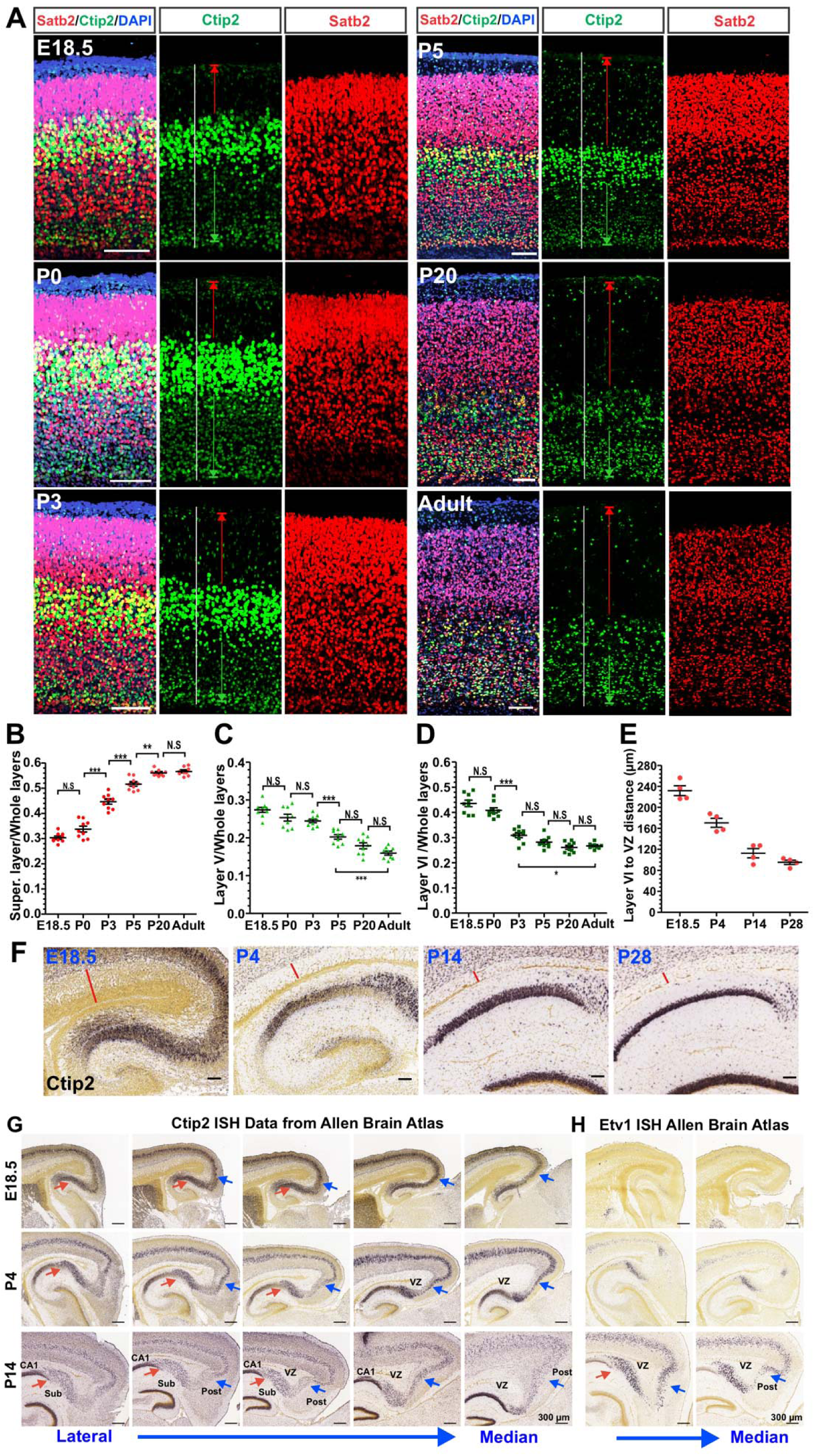
Cortical laminar changes during postnatal development indicate a slow reverse movement of neocortical neurons. **(A)** Cortical co-immunofluorescence staining using Satb2 and Ctip2 antibodies in WT mice. White line denotes whole six layers of the neocortex; red line labels the neocortical superficial sublayer starting from the top edge of Layer V to the pial surface; green line marks Layer VI starting from the bottom edge of layer V to the end of Ctip2 expression. Scale bar: 100 µm. **(B)** The ratio of superficial sublayer to whole 6 layers increased after birth and stabilized after P20. **(C)** The ratio of deep sublayer V to whole 6 layers decreased postnatally. **(D)** The ratio of Layer VI to whole 6 layers decreased significantly in the first week after birth and stabilized after P20. N = 3, mixed sexes, one-way ANOVA with Bonferroni post hoc tests. **(E)** The distances from Layer VI to the ventricular zone (VZ) decreased over ages. p < 0.0001 by one-way ANOVA. Data of **(E**) were quantified based on red line between the VZ to the edge of Layer VI using Allen Brain Atlas images **(F**). Scale bars: 100 µm. **(G-H)** Subiculum and postsubiculum exhibit a reverse movement for lamina refinement during postnatal development. **(G)** Ctip2 ISH staining from E18.5, P4 and P14 of mouse sagittal sections. Bule arrows denote the curvature between the postsubiculum that emerge at P4 and becomes remarkable at P14. Red arrows denote the demarcation between CA1 and subiculum, which becomes more obvious with age. **(H)** Similar pattern changes in the CA1, subiculum, postsubiculum and neocortex were observed in Etv1 ISH staining. Staining using other markers show similar departmental pattern. VZ: the ventricular and subventricular zone, Sub: subiculum, POST: postsubiculum. ISH Data were adopted from Allen Brain Atlas.

**Fig. S5.**
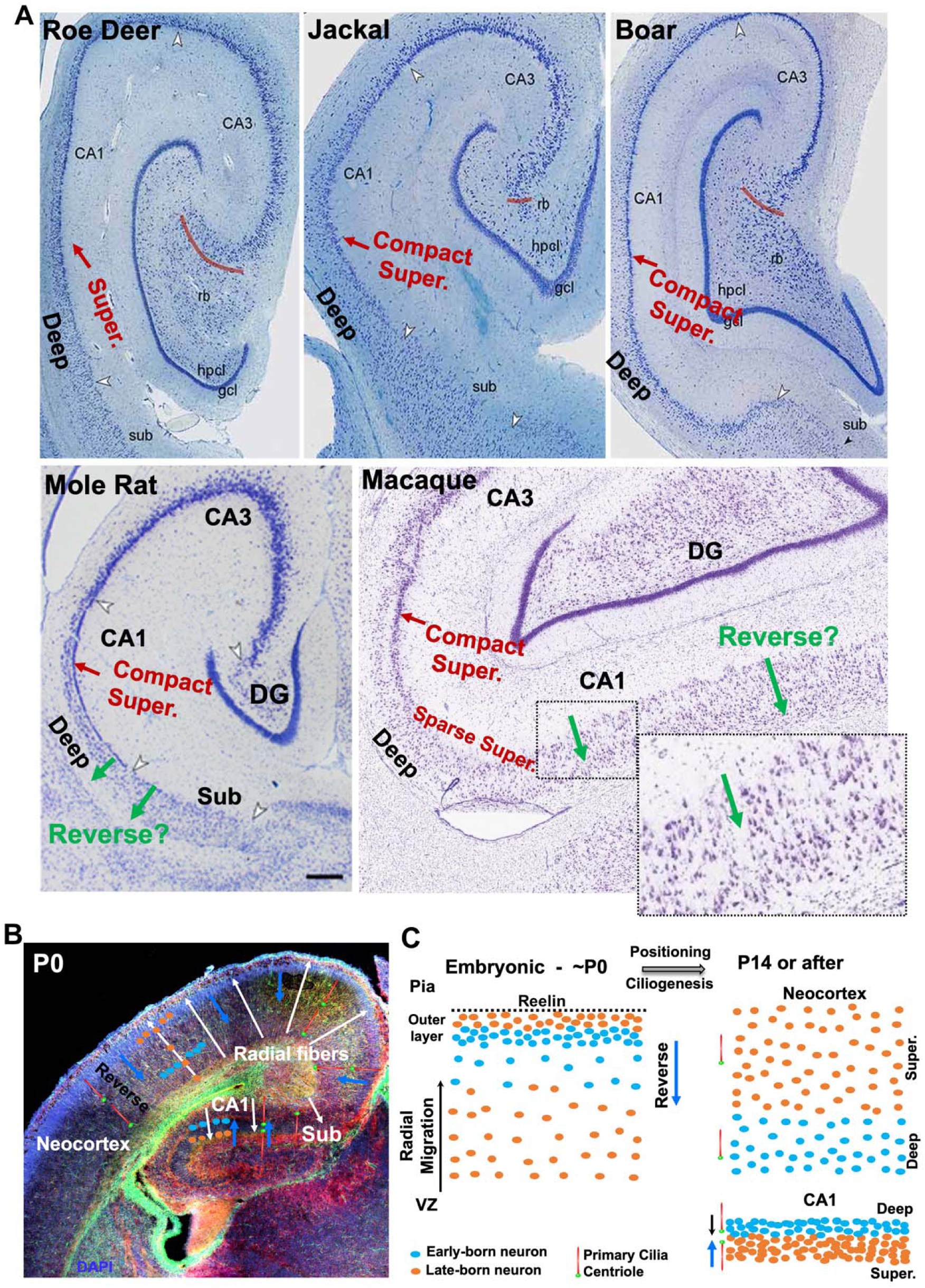
Working modeling: A slow reverse movement of principal neurons refines inside-out laminae in the cerebral cortex spanning from the hippocampal CA1 region to the neocortex. **(A)** Comparative anatomy of the mammalian hippocampi. Red arrows indicate compact superficial sublayers (Super.). Green arrows denote putative reverse direction of pyramidal neurons. Sub: subiculum. Images of roe dear, jackal, and boar are modified from (Malikovic et al., 2022), mole rat image from Radic et al., 2017, respectively. We put arrows and texts denote the direction of reverse movement. The macaque image was from NIH Non-Human Primate Brain Atlas. In macaque, the pyramidal shape of CA1 neurons is in line with putative reverse direction (enlarged on the bottom). The sparsely layered CA1 SP of primates is likely formed via reverse movement of principal neurons from the condensed superficial layer to deep sublayer. Note that the CA3 and DG principal cell layers remain highly compact. **(B)** Mouse neocortex and hippocampus at P0. After radial migration, principal neurons are subject to postnatal positioning. **(C)** A model of reverse movement for principal neuron positioning. In the neocortex and neighboring regions (including the entorhinal and cingulate cortex), both early- and late-born neurons reverse. In the compact hippocampal CA1 SP or other 3-layered allocortices, some neurons (primarily late-born neurons) reverse, whereas other neurons (mostly early-born neurons) do not. Primary cilia of principal neurons indicate the direction of reverse movement. White and blue arrows indicate the direction of radial migration of principal cells (mostly during embryonic stage) and postnatal reverse movement, respectively. Deep: deep sublayer. Super.: superficial sublayer. VZ: the ventricular zone. The early-born and late-born neurons in the CA3 are not well separated. Additionally, DG lamina is formed by outside-in migration pattern. Thus, CA3 and DG do not fit this model.

## Notes

### Competing Interest Statement

The authors have declared no competing interest.

### Summary of Updates

We revised Introduction and Result sections, modified Figures 1, 2, 3, 4 and 6, and updated Supplemental Figure S1-S3. We also revised figure legends.

